# Parallel Evaluation of Nucleophilic and Electrophilic Chemical Probes for Sulfenic Acid: Reactivity, Selectivity and Biocompatibility

**DOI:** 10.1101/2021.06.23.449646

**Authors:** Yunlong Shi, Kate S. Carroll

## Abstract

S-sulfenylation of cysteine thiols (Cys-SOH) is a regulatory posttranslational modification in redox signaling and an important intermediate to other cysteine chemotypes. Owing to the dual chemical nature of the sulfur in sulfenic acid, both nucleophilic and electrophilic chemical probes have been developed to react with and detect Cys-SOH; however, the efficiency of existing probes has not been evaluated in a side-by-side comparison. Here, we employ small-molecule and protein models of Cys-SOH and compare the chemical probe reactivity. These data clearly show that 1,3-diketone-based nucleophilic probes react more efficiently with sulfenic acid as compared to strained alkene/alkyne electrophilic probes. Kinetic experiments that rigorously address the selectivity of the 1,3-diketone-based probes are also reported. Consideration of these data alongside relative cellular abundance, indicates that biological electrophiles, including cyclic sulfenamides, aldehydes, disulfides and hydrogen peroxide, are not meaningful targets of 1,3-diketone-based nucleophilic probes, which still remain the most viable tools for the bioorthogonal detection of Cys-SOH.

## 1. Introduction

The cysteine residue undergoes a broad variety of oxidative post-translational modifications (OxiPTMs), which highlights its diverse functions in redox regulation and signaling, and warrant delicate methods for detection and characterization [1–3]. Among these cysteine OxiPTMs, the sulfenic acid modification is the initial product of thiol oxidation by a common reactive oxygen species, H_2_O_2_, and is pivotal to its transformation to other OxiPTMs, including the sulfinic acid, sulfonic acid, disulfide, persulfide, and thiosulfinate modifications [4]. When stabilized by local environmental factors [5], such as hydrogen bond acceptors, limited solvent access, and lack of adjacent nucleophiles (*e.g*. free thiols), cysteine sulfenic acid modification can modulate protein functions by altering their activity, stability, localization and ability to bind metal ions or other proteins [6]. Most sulfenic acids are transient intermediates to other OxiPTMs, primarily disulfide bridges that confer protein structural stability, or in some cases redox-active disulfides with regulatory functions [7]. Overall, profiling protein sulfenic acid modification has paved the way for elucidation of redox-based mechanisms in various organisms [8–16].

The short lifetime of protein sulfenic acids limits the use of antibody, crystallography, or mass spectrometry (MS)-based approaches for direct detection [5]. Indirect approaches monitoring the changes in thiol population upon reduction also struggle to recognize sulfenic acids among other OxiPTMs due to the lack of specificity of the reducing agents. To date, analyses of the cysteine “sulfenylome” are enabled almost exclusively by chemical probes that target this modification [3]. An ideal probe should react rapidly with sulfenic acids before they are sequestered by cellular nucleophiles (mainly thiols), and should exhibit strong chemo-selectivity, especially against thiols, which constitute >90% of global cellular cysteines [17]. Therefore, it is critical to develop chemical reactions with excellent bioorthogonality, a term comprising virtues of reactivity (fast reaction kinetics), selectivity (no off-target reaction) and biocompatibility (stability, permeability, toxicity, etc. suitable for applications in complex biological systems) [18].

Cysteine sulfenic acids possess both nucleophilic and electrophilic properties [5], which lead to several strategies for chemical ligations. Analogous to sulfenyl halides, the electrophilic sulfur atom in sulfenic acid is subject to nucleophilic attack by various nucleophiles, including carbanions, amines, phosphines, and thiols^5^. However, heteroatom-based nucleophiles tend to give reversible adducts with sulfenic acids, because of their weak bonds (N-S bond *D*°_298_ = 467 kJ/mol, S-S bond *D*°_298_ = 425 kJ/mol; compare to C-S bond *D*°_298_ = 712 kJ/mol) [19]. Consequently, carbon nucleophiles have long been used for covalent labeling of sulfenic acids, with dimedone, a 1,3-diketone compound, being the most prominent example [5]. Several groups including ours have developed hundreds of 1,3-diketone derivatives with various reporter tags and a wide range of reactivity profiles [20–22] (Fig. 1A and B, left). On the other hand, deprotonated sulfenic acids (sulfenate anions) display moderate to weak nucleophilicity, and react with strong electrophiles such as alkyl iodides or 4-chloro-7-nitrobenzofurazan (NBD-Cl) [23], albeit at low reaction rates [24]. Such reactions have found applications mostly in simple systems (small-molecules or purified proteins) due to significant cross-reactivity under a highly nucleophilic biological environment. Sulfenic acids are also known to react with alkenes and alkynes under heat [5], via a concerted cycloaddition mechanism (Fig. 1A and B, right). Ring strain-activated alkenes/alkynes have been used to promote reactivity for bioconjugation to sulfenic acids. Notable examples include a cyclooctyne reagent [25] originally used for strain-promoted azide-alkyne cycloaddition (SPAAC), and a trans-cycloctene derivative [26] originally used for tetrazine ligation. Norbornadiene also reacts with small-molecule sulfenic acids [27]; however, recent reports claimed that a less strained norbornene derivative could capture sulfenic acids as well [28].

**Fig. 1.**
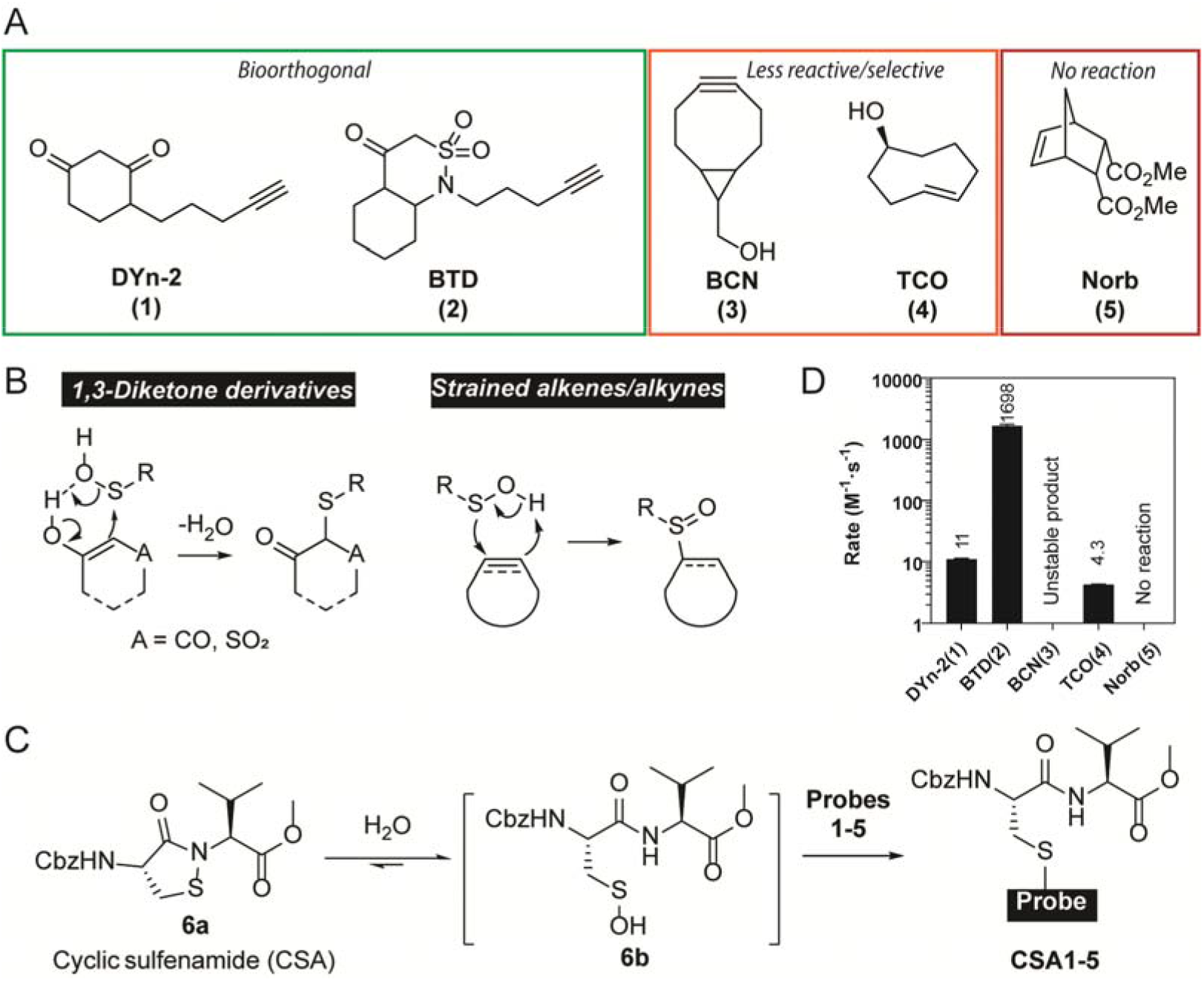
Kinetic rate studies of sulfenic acid probes with a cyclic sulfenamide (CSA) model. A) Structures of chemical probes involved in this study. B) Two types of sulfenic acid probes and their mechanisms of action. C) CSA sulfenic acid model. D) Kinetic rate of probes 1-5 reacting with CSA.

The structure-activity relationship of 1,3-diketone derived C-nucleophile probes has been well-studied [21]. Generally speaking, destabilization of the carbanion at C-2 is linked to increased reactivity. The strength of the electron-withdrawing groups (inductive effect) is the primary factor, while resonance and steric effects also play a part. In contrast, kinetic profiles of strained alkene/alkyne-based sulfenic acid probes are inconclusive, because different small-molecule or protein models were used to evaluate reaction rates. In addition, questions and conjectures with respect to the cross-reactivity of C-nucleophile probes remain unanswered within a physiological context. Here, we implement multiple models to benchmark the reactivity of various types of sulfenic acid probes, and address the selectivity of C-nucleophiles with both the rate and occurrence of potential off-targets in mind.

## 2. Results and discussion

### 2.1 Comparison of probe reactivity using small-molecule sulfenic acid models

Several recently reported probes with applications in proteome-wide detection of protein S-sulfenylation, including two 1,3-diketone derivatives DYn-2 (**1**) [8], BTD (**2**) [22], and three strained alkyne/alkene probes BCN (**3**) [25], TCO (**4**) [26], Norb (**5**) [27] were selected for our study (Fig. 1A). First, we used a dipeptide cyclic sulfenamide (CSA) **6** as a small-molecule model for sulfenic acid (Fig. 1C and D). Previously we reported that CSA readily hydrolyzes in aqueous solution to generate a transient sulfenic acid, which eventually becomes thiosulfenate and sulfinic acid in absence of chemical probes^21^. CSA has the advantages of representing a chemically unperturbed cysteine sulfenic acid in aqueous solutions, and easy handling due to its stability in organic solvents. It has been successfully used for the benchmark of a library of >100 C-nucleophiles, whose reaction kinetics with CSA correlate well with the labeling efficiency in sulfenylated proteins, cell lysates and living cells [22,29]. Furthermore, CSA kinetics are consistent with the number of MS-identified S-sulfenylation sites from the human proteome. For example, DYn-2 (**1**), which is an alkyne-functionalized nucleophilic probe based on dimedone, posed moderate reactivity with CSA at a rate of 10 M^-1^·s^-1^ and labeled 183 sites in RKO cell lysates, while BTD (**2**) is a recent probe with a vastly superior kinetic rate at 1,700 M^-1^·s^-1^ and labeled 1186 sites in the same study [22]. The cyclooctyne derivative BCN (**3**) also reacted in the CSA model, but the product found by LC-MS, presumably from the alkyne attacking the electrophilic sulfur, was not stable and eventually converted to the disulfide of the dipeptide; the desired labeling product was only observed in trace quantity. In a two-step process, the trans-cyclooctene TCO (**4**) attacks the electrophilic sulfur with its alkene moiety, followed by participation of a neighboring axial hydroxyl group. It registered a slightly slower rate than DYn-2 at 4.3 M^-1^·s^-1^. It is worth noting that unlike the 1,3-diketone derivatives, TCO functions at acidic conditions (*e.g.,* 25% formic acid) because the alkene nucleophilicity is less affected by pH changes. On the other hand, Norb (**5**) apparently did not react in the CSA model as no adduct was observed. The results above indicate that the alkene nucleophile TCO is less reactive than C-nucleophile probes, while the cyclooctyne and norbornene derivatives are not compatible with the CSA model, likely due to their intrinsic reactivity towards the sulfenamide moiety.

To develop a practical small-molecule sulfenic acid model with broad compatibility, we utilized a caged sulfenic acid precursor tagged with a fluorophore (Fig. 2A). Elimination of a sulfoxide produces a sulfenic acid and an alkene, and the rate of this reaction can be modulated by the strength of the electron-withdrawing groups on the β-carbon [30]. With this in mind, we synthesized DMDE (**7**), a BODIPY-tagged, cysteine-derived sulfoxide, which slowly releases a sulfenic acid and an alkene **8** over 16 h at 37 °C. DMDE was incubated with 10 equivalents of sulfenic acid probe **1**-**5**, then the compositions of the reaction mixtures were analyzed by HPLC (493 nm absorption for the BODIPY tag) after completion. Each mixture was composed of probe-labeled product **9a**-**e**, together with disulfide **9f** and sulfinic acid **9g** as byproducts. Similar to the prior findings with the CSA model, 1,3-diketone derivatives captured more than half of the BODIPY-tagged cysteine sulfenic acids, with BTD achieving greater than 80% yield. In comparison, BCN and TCO labeled 30% and 20% of sulfenic acids, respectively, while the rest degraded to disulfides and sulfinic acids over time. In contrast, Norb did not give any detectable labeling product, casting doubts on its ability to capture transient sulfenic acids (Fig. 2B).

**Fig. 2.**
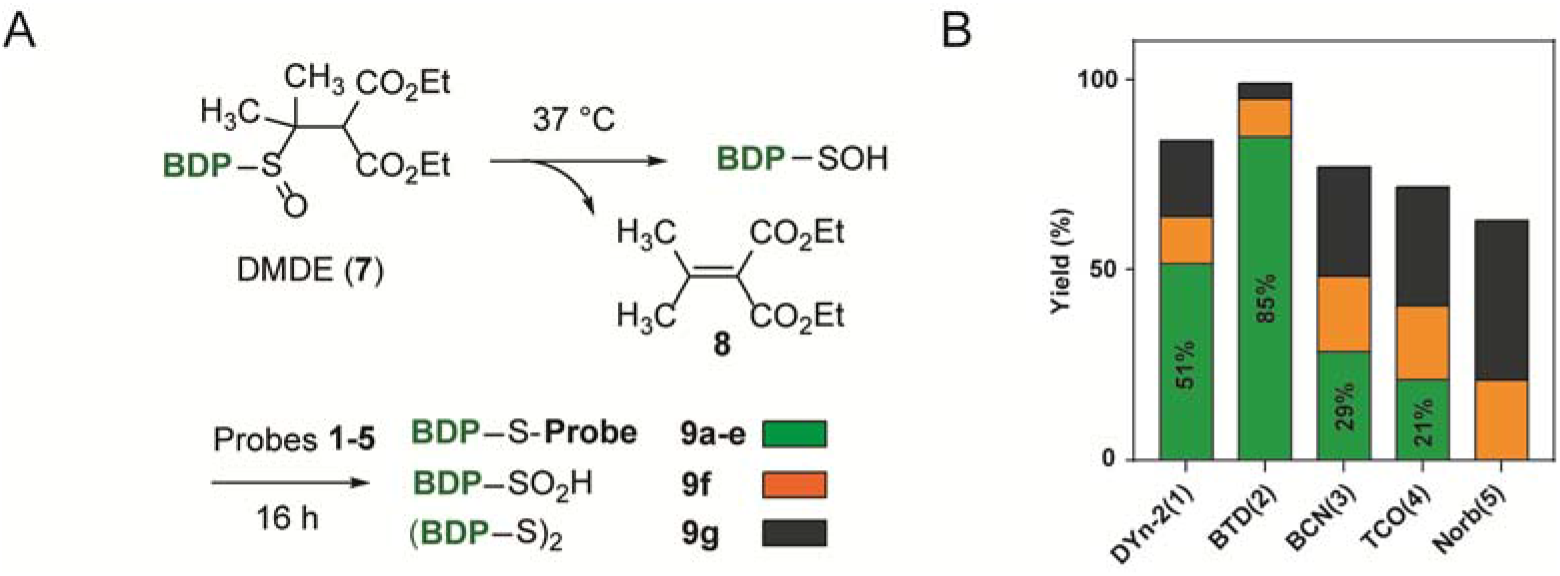
Yields of sulfenic acid capture from a fluorescent sulfoxide model. A) Elimination of a sulfoxide generated a transient sulfenic acid that was captured by probes. BDP, BODIPY-FL tag, see supplementary materials for full structures. B) Yield of captured sulfenic acids (probe adducts, green) and escaped side products (sulfinic acids in orange; disulfide in black).

### 2.2 Comparison of probe reactivity using Gpx3 protein sulfenic acid

To field-test the aforementioned probes with their intended targets, a stabilized protein sulfenic acid in C64,82S glutathione peroxidase 3 (Gpx3) was employed (Fig. 3A). This Gpx3 mutant hosts one redox-sensitive cysteine at C36 that can be readily oxidized to a sulfenic acid by a stoichiometric amount of hydrogen peroxide (H_2_O_2_) and has been successfully validated in numerous protein sulfenic acid studies [20,21,24]. We observed a similar trend in protein sulfenic acid labeling: DYn-2 captured 49% and 67% of Gpx3 (10 μM) at 0.1 mM and 1 mM (10 and 100 equivalents compared to the protein), respectively (Fig. 3B and C). BTD was the most efficient probe that quantitatively captured Gpx3 when administrated at 0.1 mM (Fig. 3D). BCN labeled just 25% of Gpx3 at a lower concentration of 0.1 mM, but the yield increased to 80% at 1 mM (Fig. 3E and F). TCO labeled 53% and 64% of Gpx3 at 0.1 and 1 mM, respectively (Fig. 3G and H). Again, no labeling product was observed with up to 100 equivalents of Norb (Fig. 3I).

**Fig. 3.**
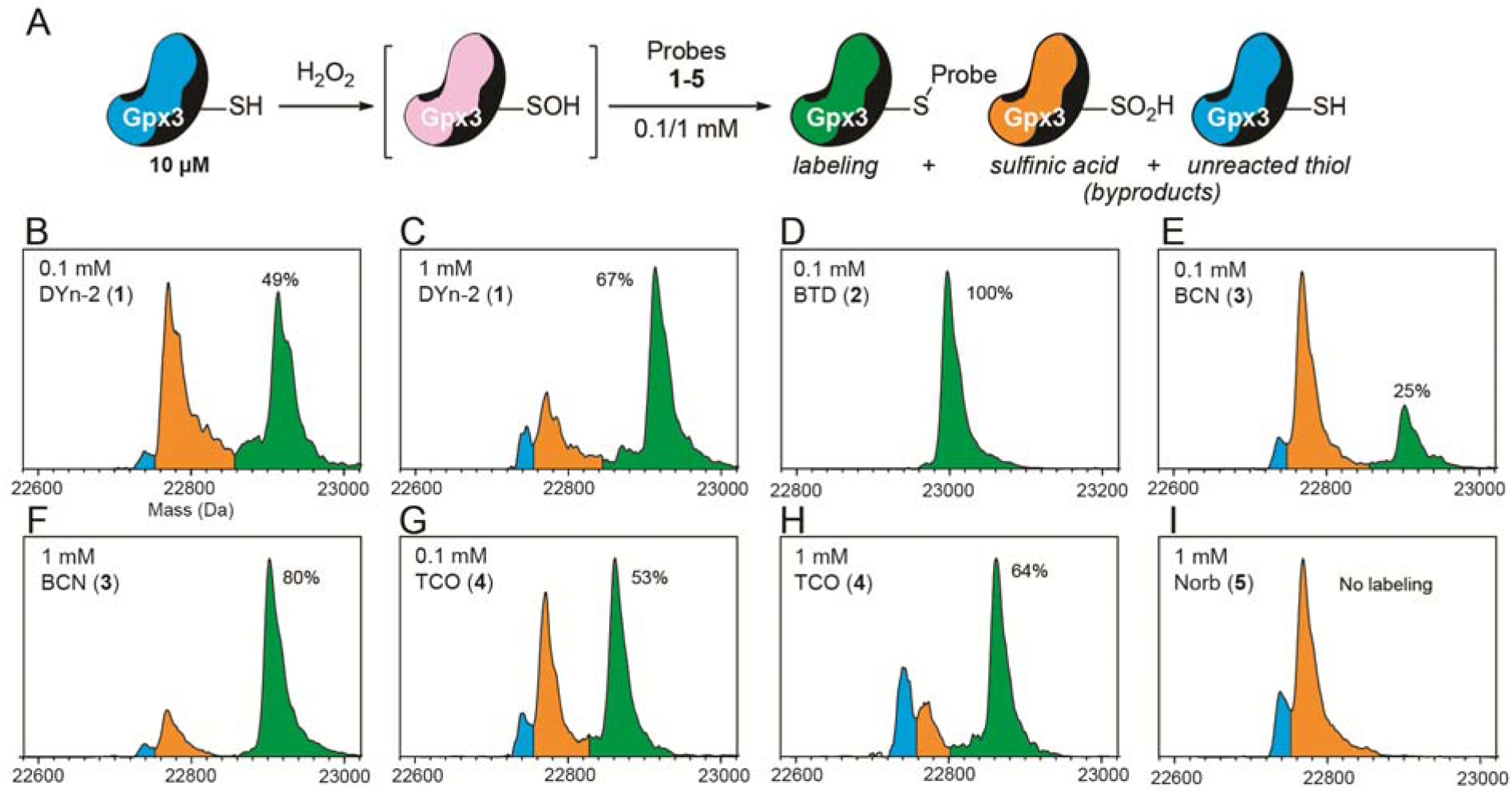
Reaction of protein Gpx3 sulfenic acid with chemical probes. A) In situ oxidation of Gpx3 (C64,82S) with quantitative H_2_O_2_ resulted in a sulfenic acid that was labeled by probes (green). Gpx3 sulfinic acid (orange) and the unreacted thiol form (blue) were also observed. B-I) Deconvoluted mass spectra of Gpx3 labeling by DYn-2 (B-C), BTD (D), BCN (E-F), TCO (G-H) and Norb (I).

In summary, the trend in reactivity of surveyed probes was BTD?DYn-2>BCN≈TCO, whereas Norb was incapable of targeting sulfenic acids in all models tested. Generally speaking, 1,3-diketone derived C-nucleophiles were more potent than strained alkynes/alkenes, whose efficacy was mainly dictated by the ring strain. This reinforces the idea that sulfenic acids reacts more efficiently with more localized electronegative charges, or “harder” nucleophiles, by the decreasing order of carbanions (*e.g.,* BTD), enolates (*e.g.,* DYn-2), strained alkynes (*e.g.,* BCN) or alkenes (*e.g.,* TCO). Norbornenes (*e.g.,* Norb) simply cannot provide enough ring strain to facilitate the reaction with sulfenic acids at a useful rate.

### 2.3 Selectivity of 1,3-diketone-based nucleophilic probes in biological context

We have demonstrated that exploiting the electrophilic property of sulfenic acids with nucleophilic probes is the preferred strategy for labeling. Although the cellular environment is also nucleophilic as a whole, it is critical to carefully evaluate the potential cross-reactivity with other biologically relevant electrophiles. Recently, such concerns were raised on several species, including the cyclic sulfenamides, aldehydes, disulfides and hydrogen peroxide, but these analyses were mostly based on isolated chemical models without biological contexts. Therefore, we move to clarify these issues with protein models and kinetic studies.

Cysteine sulfenamides are formed via an attack of an amide nitrogen on a sulfenic acid. Favored by conformation, the amide is from the backbone of the following amino acid in the sequence, yielding a five-membered cyclic structure. Like sulfenic acids, some cyclic sulfenamides can be good electrophiles and produce the same product when reacting with C-nucleophiles. For example, the dipeptide sulfenamide CSA can react with several thiol, phosphinate, and carbon nucleophiles in an anhydrous environment [31]. However, the formation of a (cyclic) sulfenamide will face a huge hurdle, because under normal circumstances, an amide (p*K*_a_ ~ 17) is a very poor nucleophile and a sulfenic acid has a very short lifetime. Although the sulfenamide modification is rarely observed in proteins, a cyclic sulfenamide structure in protein tyrosine phosphatase 1B (PTP1B) was characterized crystallographically [32,33], which draws questions as to whether the target of 1,3-diketone derived C-nucleophiles is actually the sulfenamide [34,35]. In fact, the formation of PTP1B sulfenamide is facilitated by two key factors, including a decreased p*K*_a_ (increased nucleophilicity) of the Ser215 amide nitrogen due to hydrogen bonding, and steric shielding of the Cys216 sulfenic acid which becomes inaccessible by small-molecule nucleophiles, *e.g.,* dimedone [36]. To analyze the reactivity of PTP1B, we added DYn-2 or BTD to PTP1B, which was either oxidized in situ, or pre-oxidized prior to nucleophile treatment, representing the sulfenic acid (-SOH) or sulfenamide (-SN) form of PTP1B, respectively (Fig. 4A). Neither nucleophile reacted with PTP1B-SN, while only a small percentage (10%) of PTP1B-SOH reacted with BTD. A PTP1B reaction-based inhibitor (RBI) **10** [37] which labeled the majority (70%) of PTP1B-SOH also failed to react with the PTP1B-SN form. These findings further indicate that the pocket harboring PTP1B-SN is not exposed for nucleophilic attack, regardless of the probe’s reactivity towards sulfenic acids. Besides PTP1B, there are only a few reports on protein cyclic sulfenamides, found in a purified protein [38] or in the gas phase of MS [25], yet not known to be relevant in cellular environments.

**Fig. 4.**
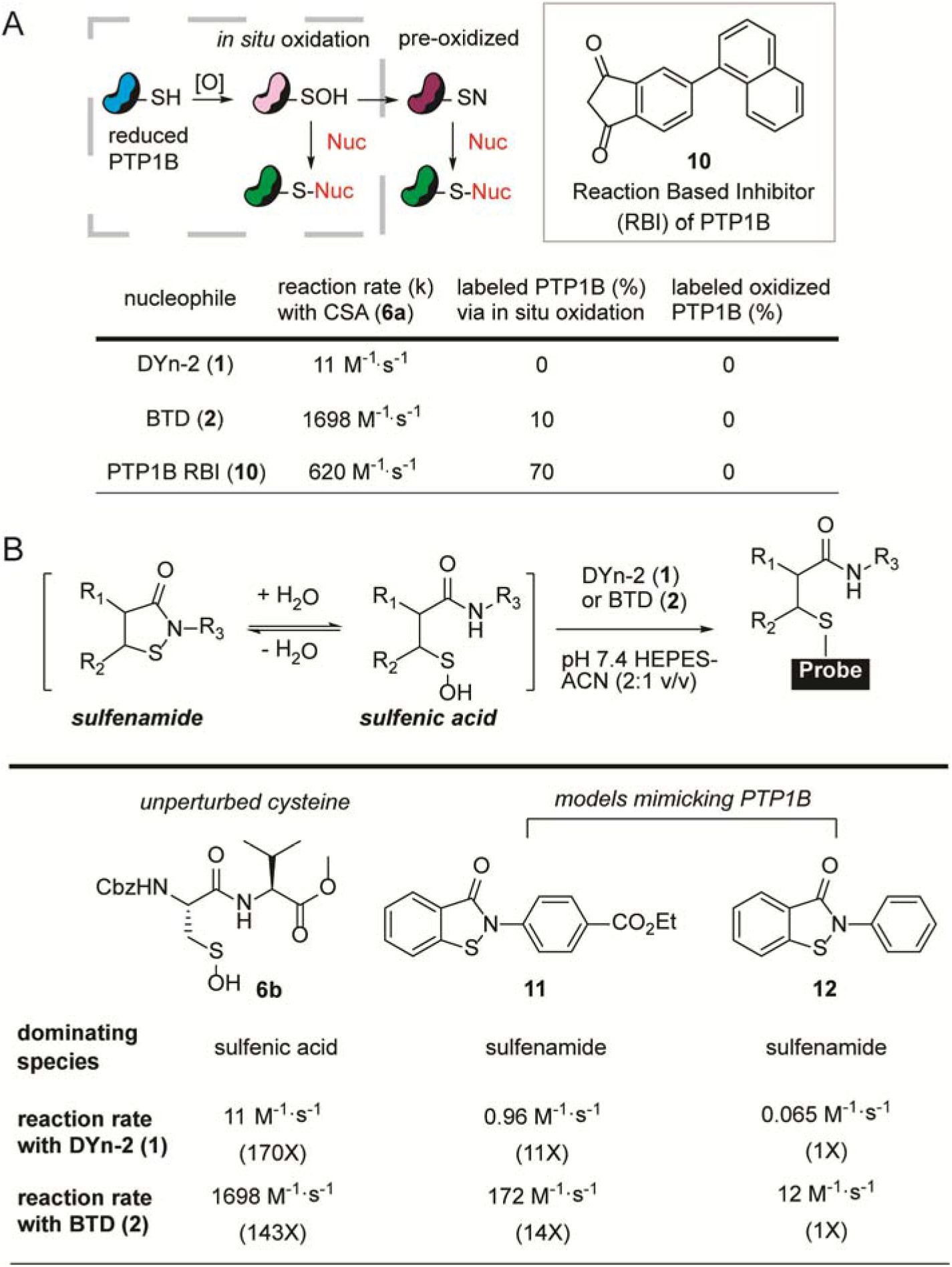
Reaction of cyclic sulfenamides with nucleophiles or probes. A) In situ oxidation and pre-oxidation of PTP1B furnished the sulfenic acid (-SOH) and sulfenamide form (-SN), respectively. The latter was inaccessible to nucleophiles including a reaction based inhibitor (RBI) of PTP1B. B) Equilibrium between a sulfenamide and a sulfenic acid. Unperturbed cysteines and synthetic models of PTP1B exhibited distinct behaviors.

In addition to the rarity in the proteome, nucleophilic attack on a sulfenamide is kinetically unfavorable and is easily outcompeted by a sulfenic acid [3]. We employed two small-molecule sulfenamide models, benzisothiazolinone derivatives **11** and **12**, which include a low p*K*_a_ thiol (~5.6, similar to the Cys216 of PTP1B [39]) and an activated amide nitrogen mimicking PTP1B. Their sulfenamide forms are the dominant species in aqueous-organic buffers, and they showed a significant decrease in reaction rates toward nucleophiles such as DYn-2 or BTD, compared to the unperturbed cysteine dipeptide **6b** (Fig. 4B). The less electron-withdrawing sulfenamide **12** suffered a greater rate decrease (>100-fold) than sulfenamide **11** (>10 fold). Thus, current evidence does not support that cyclic sulfenamide is a universal protein cysteine modification, nor a meaningful target of C-nucleophile probes.

Next, we investigated the C-nucleophile probes’ reactivity toward disulfides, which has redox potentials (*E*°) from −95 to −470 mV [40] that translate to a wide range of electrophilicity. We selected glutathione disulfide (GSSG) and a protected cystine (Z-Cys-OH)_2_ to represent redox-active disulfides which have similar redox potentials to classic oxidoreductases like thioredoxin-1 (*E*° = −270 mV) [41]. We also included a highly polarized disulfide known as 4,4′-dipyridyl disulfide (4-DPS). This electrophilic disulfide is not present in cells but is useful to model the upper limit of nucleophilic probe reactivity. Consistent with our earlier findings [21], both DYn-2 and BTD remain inactive toward GSSG or (Z-Cys-OH)_2_ (10 equivalents). Meanwhile, a rather sluggish but measurable reaction of BTD with 4-DPS (10 equivalents) was obtained (0.159 M^-^ 1·s^-1^), which is about 10,000 times slower than BTD reacting with the CSA sulfenic acid model (1,700 M^-1^·s^-1^). A small portion of DYn-2 also reacted with 4-DPS, but the reaction remained incomplete after 24 h. These data indicate that 1,3-diketones and protein disulfides are highly unlikely to be biologically relevant reaction partners (Fig. 5A and C).

**Fig. 5.**
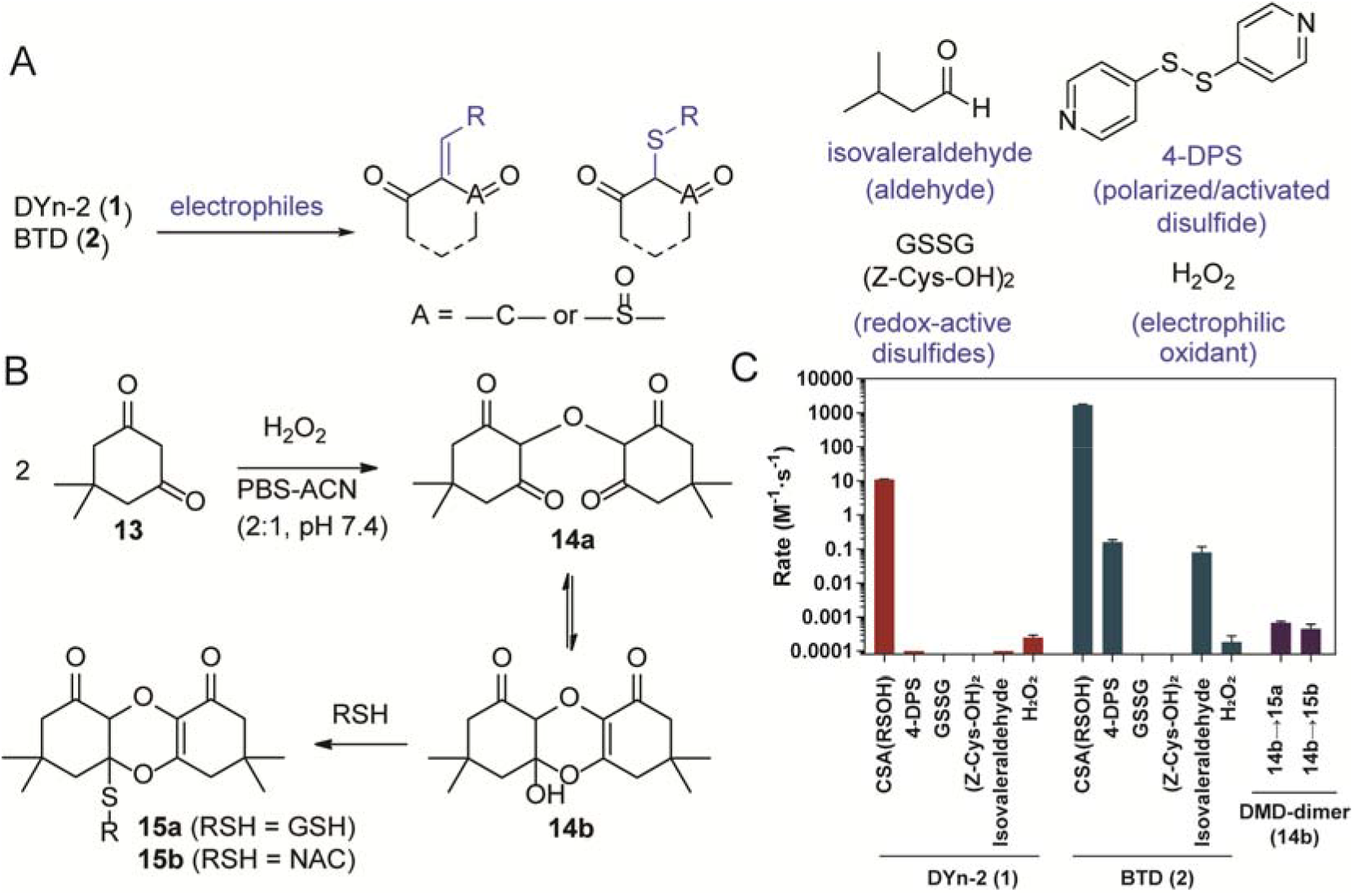
Reaction of nucleophilic sulfenic acid probes with electrophiles. A) 1,3-Diketone derived probes were reacted with various electrophiles. B) Dimerization of dimedone and the following reaction with thiols. C) Rate constants of probes reacting with electrophiles, including sulfenic acids (intended targets), aldehydes, disulfides and hydrogen peroxide (cross-reactions).

Another noteworthy class of biological electrophiles are aldehydes. They are mainly present at low levels as small-molecule byproducts of cellular metabolism [42], but some exist as modifications to biomolecules, such as the N-formylation of proteins [43]. We chose isovaleraldehyde, which possesses an alkyl side chain, to represent biologically relevant aldehydes. Similar to the reaction with 4-DPS, BTD presented poor reactivity with isovaleraldehyde at a rate of 0.079 M^-1^·s^-1^, about 21,000-fold less than that of CSA sulfenic acid model. The rate of DYn-2 and isovaleraldehyde was also too slow to be measured before aerobic degradation took place. With the kinetic data and concentration in mind, aldehydes also pose limited impact on the labeling of protein sulfenic acids by 1,3-diketone derivatives (Fig. 5A and C).

Last but not least, we address the potential cross-reactivity with a ubiquitous reactive oxygen species, hydrogen peroxide (H_2_O_2_), which is also an electrophile. It has been reported that dimedone underwent oxidative dimerization when exposed to excessive (high millimolar) concentrations of H_2_O_2_ [44]. However, the impact of this reaction is unclear under practical labeling conditions. When 5 mM of dimedone-based probe DYn-2 (typical concentration used in lysate or cell labeling experiments) was treated with 10 mM of H_2_O_2_, less than 5% of oxidative dimer was found after 2 h at 37 °C (typical labeling condition). Even after overnight incubation, DYn-2 dimerization was only about 50% complete. When H_2_O_2_ concentration was further decreased to 1 mM, which is still a few orders of magnitude higher than the estimated cellular level (low micromolar range), almost no dimer was observed after 2 h, and about 12% of DYn-2 dimerized overnight. This indicates cellular H_2_O_2_ does not degrade DYn-2 at an observable level. The more reactive nucleophilic probe BTD was also surveyed, and found to react extremely slowly with H_2_O_2_ at a rate of 1.83×10^-4^ M^-1^·s^-1^, similar to that of DYn-2 (2.48×10^-4^ M^-1^·s^-1^) (Fig. 5B and C).

The oxidative dimerization of dimedone (**13**) was further investigated by NMR. Data showed the dimer existed as a mixture of enol-keto isomers in chloroform. As expected, the equilibrium shifted to the keto form in polar solvents like DMSO, yielding two singlets with a 2:3 ratio in ^1^H NMR, which suggested a symmetric structure **14a**. We also observed that **14a** reacted with thiols like GSH and NAC, likely through a hemiketal intermediate **14b** and a thiol-acetal exchange reaction to furnish **15a** and **15b**. Nevertheless, these reactions occurred at decreased rates (6.81×10^-4^ M^-1^·s^-1^ and 4.42×10^-4^ M^-1^·s^-1^, respectively) (Fig. 5B and C). Taken together with the already sluggish dimerization reaction, cellular levels of H_2_O_2_ will not interfere with the specificity and efficacy of 1,3-diketone-based sulfenic acid probes, and dimedone is not expected to form covalent bonds with thiols via the oxidized dimer under physiological settings.

## 3. Conclusion

Reactivity and selectivity are the most critical features of chemical probes for bioorthogonal applications. With the recent emergence of new classes of sulfenic acid probes, we evaluated them in parallel using two small-molecule models with focus on their kinetic rates or reaction yields, and further validated the trend in reactivity using a protein model. In contrast to 1,3-diketone based nucleophiles (*e.g.*, DYn-2 (**1**), BTD (**2**)), some strained alkene/alkyne derivatives (*e.g.,* BCN-OH (**3**), TCO-OH (**4**)) target sulfenic acids via different mechanisms, but at a lower efficacy even compared to the first-generation dimedone-based nucleophilic probes. We found no evidence of the norbornene derivative Norb (**5**) reacting with all three of our sulfenic acid models. The striking difference from the original reports are most likely attributed to an unsound model, which utilizes excessively high concentrations of *N*-acetylcysteine and H_2_O_2_ ostensibly to generate sulfenic acid, but instead yield thiyl radicals which are sequestered by norbornenes.

Aerobic oxidation of thiols can also generate thiyl radical, which are chief participants in thiol-ene addition reactions [45] with strained alkene/alkyne probes. On the other hand, concerns have been expressed regarding the potential for cross-reactivity of nucleophilic probes with electrophiles; however, the population is such species is limited by the overall nucleophilic cellular environment. Our results indicate that common bioelectrophiles – aldehydes, polarized (activated) disulfides most prone to nucleophilic attack, and electrophilic oxidant (H_2_O_2_), react with 1,3-diketone-based probes at rates several orders of magnitude less than the intended target, sulfenic acids. Additionally, we provide a detailed analysis of cyclic sulfenamide reactivity, formation and abundance, which indicates that such species are extremely rare in biology. Two benzisothiazolinone derivatives that mimic the Ser215-Cys216 motif of PTP1B, spontaneously formed cyclic sulfenamides *via* sulfenic acid intermediates, but showed reduced reactivity toward nucleophiles compared to an unperturbed cysteine. These findings, with steric considerations, illustrates why most nucleophiles, including inhibitors designed to fit the active site, fail to target the PTP1B sulfenamide.

Although small-molecule models are versatile tools for the generation of kinetic data, they are generally not soluble in purely aqueous solvent and addition of organics (to facilitate solubilization) can have a dramatic impact on reaction rates as well as preferential stabilization of cysteine oxoforms. For example, solvents dictate the dominating species of synthetic sulfenamides like CSA, which exists in stable cyclized form (**6a**) in organic solvents, but rapidly hydrolyzes to the sulfenic acid form (**6b**) in aqueous solutions. As another example, anthraquinone-1-sulfenic acid (Fries’ Acid) is stabilized *via* an intramolecular hydrogen bond. It exhibits a vastly different reactivity profile compared to protein sulfenic acids and is hence not an ideal model for the latter [25]. Protein sulfenic acids models may also exhibit different rates depending upon factors such as solvent accessibility, intrinsic sulfur reactivity, and the presence of acidic or basic amino acids. This limitation is underscored by the reaction of oxidized PTP1B, whose buried sulfenamide does not react with most nucleophiles, while positive reactivity is observed in small-molecule “models” of the PTP1B sulfenamide. Therefore, it is critical to establish standardized (preferably, protein-based) models for testing new probes for cysteine oxoforms. The present study evaluates nucleophilic/electrophilic probe reactivity in both protein and small-molecule sulfenic acid models. In conclusion, chemical probes based on the 1,3-diketone scaffold remain the most viable bioorthogonal approach to trapping and tagging protein sulfenic acids.

## 4. Methods

All associated methods are included in the supplementary materials.

## Acknowledgements

This work was supported by the US National Institutes of Health (R01 GM102187 and R01 CA174864 to K.S.C).

