## supplementary materials for "Parallel Evaluation of Nucleophilic and Electrophilic Chemical Probes for Sulfenic Acid: Reactivity, Selectivity and Biocompatibility"

**Appendix A. Supplementary Material**

Yunlong Shi and Kate S. Carroll*

*

### SI1. Materials and general synthetic methods

All reactions were conducted in flame-dried glassware under nitrogen pressure with dry solvents, unless otherwise noted. All reagents and solvents were purchased from Sigma-Aldrich (St. Louis, MO) or Fisher Scientific (Waltham, MA) and used as received. Reactions were monitored by thin layer chromatography (TLC) carried out using MilliporeSigma TLC silica gel 60 plates with F254 coating. Crude reaction mixtures were purified using a CombiFlash Nextgen 300+ chromatography system with prepacked silica gel columns. ^1^H NMR and ^13^C NMR spectra were collected in CDCl3 or DMSO-d_6_ (Cambridge Isotope Laboratories, Cambridge, MA) at 400 and 100 MHz respectively, using a Bruker AM-400 instrument with chemical shifts relative to residual CHCl3 (7.27 and 77.37 ppm) or DMSO (2.50 and 39.52 ppm). Low resolution mass spectra were acquired from an Agilent 6120 quadruple LC/MS system. All spectra are available upon request.

### SI2. Synthetic procedures

DYn-2 (**1**), BTD (**2**) and CSA (**6a**) were prepared according to literature procedures^2-4^. Compound **3-5** were purchased from commercial sources (see above) and used as received.

**Synthetic procedure for the redox-caged sulfenic acid precursor, DMDE (7)**

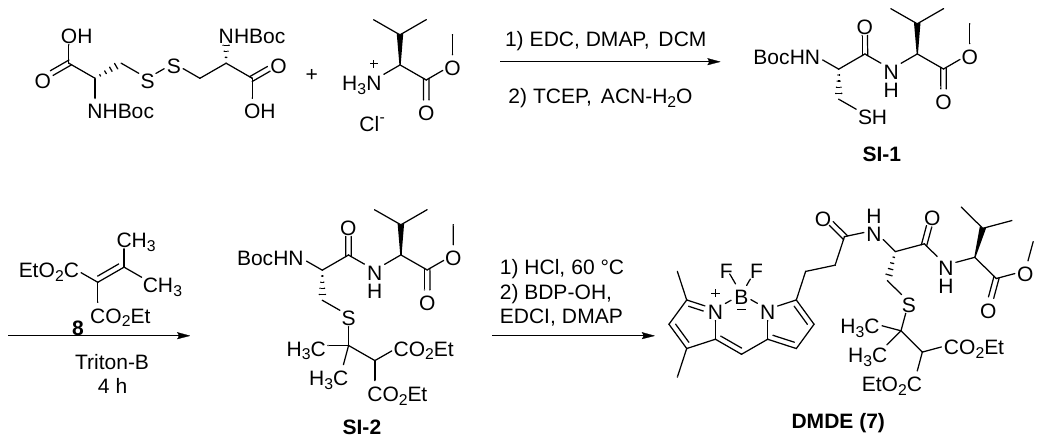

**Preparation of the Boc-Cys-Val-OMe dipeptide (SI-1)**

To a flask containing (Boc-Cys-OH)_2_ (440.5 mg, 1.0 mmol), *L*-Valine methyl ester hydrochloride (502.9 mg, 3.0 mmol) in 20 ml dry DCM at 0 °C was added EDCI⋅HCl (575.1 mg, 3.0 mmol) and DMAP (12.2 mg, 0.1 mmol). The mixture was slowly warmed and stirred at room temperature overnight, diluted with EtOAc (60 ml) washed with sat’d NaHCO_3_ (30 ml x 2) and brine (30 ml). The organic layer was dried, concentrated and purified by chromatography (20-50% EtOAc in hexane) to afford the disulfide (Boc-Cys-Val-OMe)_2_ as a white solid, which was dissolved in ACN-H_2_O (15 ml + 15 ml) was added tris(2-carboxyethyl)phosphine hydrochloride (TCEP⋅HCl, 573.3 mg, 2.0 mmol). The solution was stirred at 37 °C for 5 h. After completion, it was diluted in DCM (20 ml) and washed with sat’d NaHCO_3_ (20 ml x 2) and brine (20 ml), dried (Na_2_SO_4_) and concentrated *in vacuo* to afford the compound **SI-1** as a white solid (185.1 mg, 0.553 mmol, 55% yield). **^1^H NMR** (400 MHz, CDCl_3_) δ 6.85 (brs, 1H), 5.48 (brs, 1H), 4.51 (dd, *J* = 8.7, 4.9 Hz, 1H), 4.34 (m, 1H), 3.73 (s, 3H), 3.05 (m, 1H), 2.73 (m, 1H), 2.19 (m, 1H), 1.69 (m, 1H), 1.45 (s, 9H), 0.92 (dd, *J* = 11.9, 6.9 Hz, 6H). **^13^C NMR** (100 MHz, CDCl_3_) δ 172.17, 170.38, 155.69, 80.83, 57.56, 55.69, 52.44, 31.28, 28.47, 26.82, 19.23, 17.92. **MS** (ESI) Calculated for C_14_H_26_N_2_NaO_5_S [M+Na^+^] 357.1; observed 357.2.

**Preparation of the Boc-protected precursor SI-2**

To a flask containing 66.8 mg (0.20 mmol) of dipeptide **SI-1** in 2 ml dry THF at -78°C was added Triton-B (8 µL, 40% in methanol, 0.02 mmol) in 2 ml dry THF dropwise, then 30.0 mg diethyl isopropylidenemalonate (compound **8**, 0.15 mmol) in 2 ml dry THF was added dropwise. The reaction mixture was then stirred at room temperature until completion of reaction (about 4 h), then diluted with brine (20 ml) and extracted with DCM (20 ml x 3). The combined organic layers were dried (Na_2_SO_4_), concentrated and purified by chromatography (20-50% EtOAc in hexane) to afford **SI-2** as a colorless oil (34.6 mg, 0.065 mmol, 43% yield). **^1^H NMR** (400 MHz, CDCl_3_) δ 6.97 (brs, 1H), 5.49 (brs, 1H), 4.47 (dd, *J* = 8.7, 4.9 Hz, 1H), 4.18 (m, 4H), 3.70 (s, 3H), 3.04 (dd, *J* = 12.9, 6.6 Hz, 1H), 2.88 (dd, *J* = 12.9, 6.1 Hz, 1H), 2.15 (m, 1H), 1.55 (s, 3H), 1.53 (s, 3H), 1.43 (s, 9H), 1.25 (td, *J* = 7.1, 2.3 Hz, 6H), 0.91 (dd, J = 8.9, 6.9 Hz, 6H). **^13^C NMR** (100 MHz, CDCl_3_) δ 171.91, 170.49, 167.36, 167.12, 80.40, 61.53, 61.45, 60.41, 57.42, 52.12, 46.06, 31.29, 30.32, 28.34, 26.88, 26.54, 19.02, 17.77, 14.11. **MS** (ESI) Calculated for C_24_H_43_N_2_O_9_S [M+H^+^] 535.3; observed 535.3.

**Preparation of DMDE (7)**

A solution of 26.7 mg **SI-2** (0.05 mmol) in 2 ml HCl (4.0 M solution in dioxane) was microwaved at 60 °C for 20 min. After cooling, the solution was concentrated *in vacuo*, taken in DCM (10 ml) and washed with sat’d NaHCO_3_ solution (10 ml). The aqueous layer was extracted with DCM (10 ml x 2). Combined organic solutions were dried (Na_2_SO_4_) and concentrated *in vacuo*. The product above and 14.6 mg 3-BODIPY-propanoic acid (BDP-OH, 0.05 mmol) was dissolved in 10 ml dry DCM at 0 °C, followed by addition of 9.6 mg EDCI·HCl (0.05 mmol) and 1.2 mg DMAP (0.01 mmol). The mixture was slowly warmed to room temperature while stirring for 3 h. After completion, the mixture was diluted with DCM (10 ml) and washed with saturated NaHCO_3_ (20 ml) and brine (20 ml). The organic layer was dried (Na_2_SO_4_), concentrated and purified by flash chromatography (10-50% EtOAc in hexane) to afford compound **7** as a dark red solid (19.8 mg, 0.028 mmol, 56% yield). **^1^H NMR** (400 MHz, CDCl_3_) δ 7.15 (d, *J* = 8.6 Hz, 1H), 7.07 (s, 1H), 6.86 (d, *J* = 4.0 Hz, 1H), 6.78 (d, *J* = 6.8 Hz, 1H), 6.28 (d, *J* = 4.1 Hz, 1H), 6.10 (s, 1H), 4.61 (td, *J* = 7.1, 5.8 Hz, 1H), 4.44 (dd, *J* = 8.6, 5.0 Hz, 1H), 4.20 (m, 4H), 3.79 (s, 1H), 3.71 (s, 3H), 3.30 (t, *J* = 7.5 Hz, 2H), 3.02 (dd, *J* = 13.6, 5.8 Hz, 1H), 2.72 (m, 3H), 2.55 (s, 3H), 2.24 (s, 3H), 2.17 (m, 1H), 1.58 (s, 3H), 1.55 (s, 3H), 1.26 (t, J = 7.1 Hz, 6H), 0.93 (t, *J* = 6.7 Hz, 6H). **^13^C NMR (100 MHz, CDCl_3_)** δ 172.28, 171.83, 170.33, 167.82, 167.31, 160.52, 157.31, 143.96, 135.32, 133.50, 128.23, 123.91, 120.55, 117.28, 61.69, 61.55, 60.48, 57.75, 53.09, 52.18, 46.33, 35.52, 31.05, 30.17, 27.05, 26.37, 24.69, 19.14, 17.87, 15.06, 14.15, 11.42. **MS** (ESI) Calculated for C_33_H_48_BF_2_N_4_O_8_S [M+H^+^] 709.3 observed 709.3.

**Synthetic procedure for compound 10**

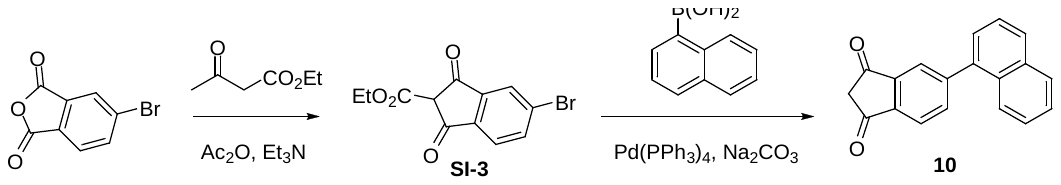

To a solution of 4-bromophthalic anhydride (10g, 44.05 mmol) dissolved in DCM (200 ml) was added ethyl acetoacetate (6.74 ml, 52.86 mmol), acetic anhydride (8.32 ml, 88.10 mmol), and triethylamine (18.42 ml, 132.15 mmol). The mixture was stirred at room temperature for 4 hours. The solution was concentrated, H_2_O was added (50 ml), followed by 2 M HCl (200 ml) at 0 °C. The yellow precipitate was filtered, washed with cold 2 M HCl, and dried. 6.93 g (53%) of the desired product (**SI-3**) was recovered and stored at 0 °C away from light.

A mixture of **SI-3** (296 mg, 1.00 mmol), 1-naphthaleneboronic Acid (344 mg, 2.00 mmol), and Na_2_CO_3_ (530 mg, 5.00 mmol) in toluene (5 ml), H_2_O (2 ml), and EtOH (1 ml) was degassed for 5 minutes with a stream of N_2_. Pd(PPh_3_)_4_ (57.7 mg, 5 mol%) was added and the mixture was heated to 100 °C for 18 hours. The reaction was cooled and filtered through celite with EtOAc washes. 2 M HCl was added and stirred for 1 hour at room temperature. The mixture was extracted (3X) with EtOAc, dried over MgSO_4_, filtered, and concentrated. The crude material was purified by flash column chromatography using a mixture of EtOAc and hexanes to afford the title compound as a pale yellow solid (202.3 mg, 0.744 mmol, 74% yield). **^1^H NMR** (400 MHz, CDCl_3_) δ 8.28 (m, 1H), 8.14 (m, 3H), 7.93 (m, 3H), 7.77 (dd, *J* = 8.5, 1.9 Hz, 1H), 7.56 (m, 2H), 3.30 (s, 2H). **^13^C NMR** (100 MHz, CDCl_3_) δ 197.76, 197.13, 148.89, 144.28, 142.05, 136.07, 134.81, 133.60, 133.47, 129.29, 128.64, 127.88, 127.22, 127.05, 124.97, 123.89, 121.54, 45.61. **MS** (ESI) Calculated for C_19_H_13_O_2_ [M+H^+^] 273.1 observed 273.1.

**Synthetic procedure for compound 14a**

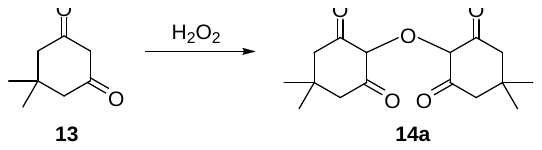

140.0 mg dimedone (**13,** 1.00 mmol) was dissolved in 10 ml Tris-HCl buffer (500 mM, pH 7.4). 1 ml hydrogen peroxide solution (30%) was added to the solution which was stirred for 3 h at room temperature. After completion, the solution was acidified with 2 M HCl to pH ~2, and extracted with EtOAc (20 ml X 3). The combined organic layers were dried (Na_2_SO_4_), concentrated *in vacuo*, and purified by flash chromatography (15-60% EtOAc in hexane) to afford compound **14a** as a pale yellow solid (92.9 mg, 0.316 mmol, 63% yield). **^1^H NMR** (400 MHz, DMSO-d_6_) δ 2.29 (s, 8H), 1.04 (s, 12H). **^13^C NMR** (100 MHz, DMSO-d_6_) δ 173.02, 44.71, 31.69, 27.16, 24.81. **MS** (ESI) Calculated for C_16_H_23_O_5_ [M+H^+^] 295.2 observed 295.3.

### SI3. Instrumentation and calculation for the LC-MS kinetic assay

Agilent technologies (Santa Clara, CA) 1220 Infinity LC coupled with a 6120 quadrupole MS system was used in the kinetic assay. Separation was performed with an Agilent Poroshell 120 SB C-18 (3.0 x 50 mm, 2.7 µM particle size) column running a gradient (5-100% H_2_O in acetonitrile) of LC-MS grade solvents with 0.1% formic acid. Data was acquired by the LC/MSD ChemStation software (Rev. B.04.03-SP1).

First order rate constant (*k*) was obtained by plotting the UV peak area (*Y*-axis) of the product against time (*X*-axis) and analyzing the plot using the “Dissociation - One phase exponential decay” function in GraphPad Prism 7:

$$Y=\left( Y0-NS \right)\cdot e^{-kX}+NS$$

Second-order rate constants (*K*) were calculated from first order rate constants (*k*) and the concentration of probes (*C*):

$$K= \frac{k}{C}$$

### SI4. Pseudo first order rate plots of CSA (6a) and probes 1, 2 and 4

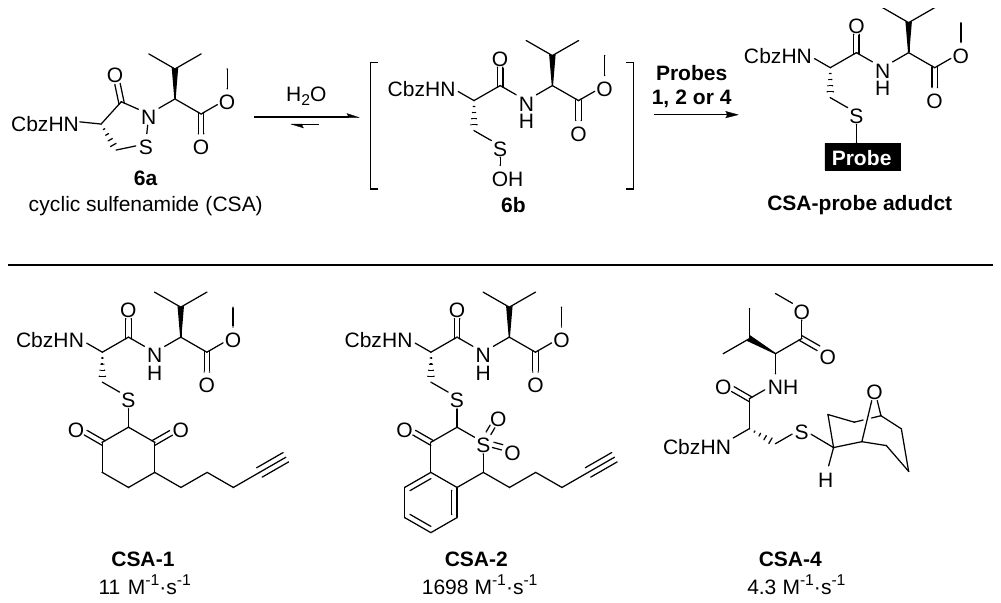

The rate studies measuring the reaction between nucleophiles with a model dipeptide-SOH were performed as previously described (*Chem. Sci.*, **2016**, *7*, 400–415). To a 2 ml solution of a nucleophile in PBS buffer (10 mM PBS pH 7.4) was added 1 ml solution of cyclic sulfenamide (CSA) in acetonitrile. In order to obtain pseudo first-order kinetics, the concentration of nucleophile was maintained at least 5 times higher than that of CSA. An aliquot (300 μl) of the reaction was collected at regular intervals (a total of 9 data points) and immediately quenched by addition of formic acid (100 μl). For the probe TCO (**4**), the reaction was quenched with 0.5 mM of 1-(triphenylphosphoranylidene)-2-propanone, which immediately scavenged the remaining CSA in solution.

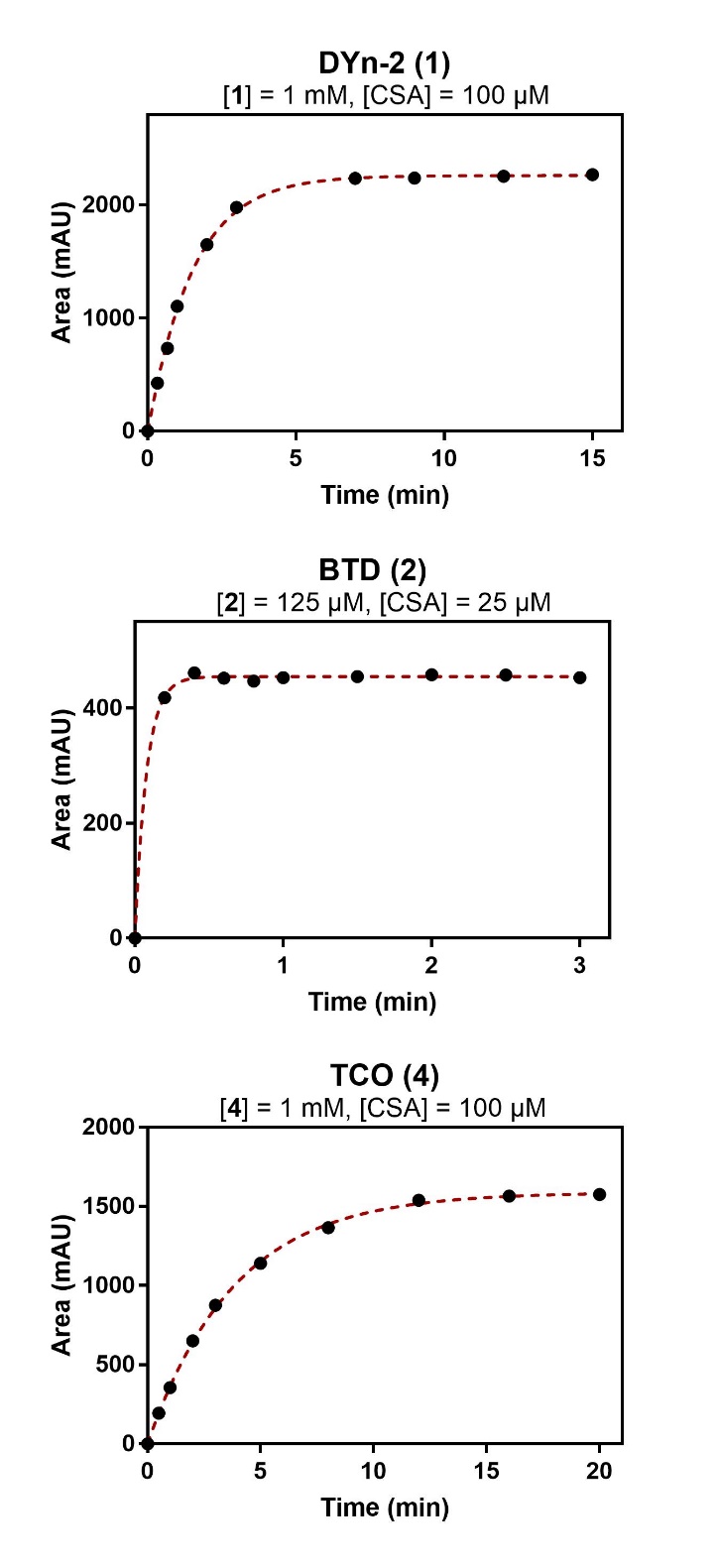

Figure S1. First order kinetic plots of probes **1**, **2** and **4** reacting with CSA (**6a**).

Table S1. Reaction table and calculations of probes **1**-**5** reacting with CSA (**6a**).

| Probe | Probe conc. | CSA sulfenic acid model (6a) conc. | First order rate, *k* (min^-1^) | Calc’d 2^nd^ order rate, *K* (M^-1^·s^-1^) |
| --- | --- | --- | --- | --- |
| DYn-2 (1) | 1 mM | 100 µM | 0.659 | 11.0 |
| BTD (2) | 125 µM | 25 µM | 12.74 | 1698 |
| BCN (3) | 1 mM | 100 µM | Unstable labeling product (see below) | |
| TCO (4) | 1 mM | 100 µM | 0.2582 | 4.303 |
| Norb (5) | 1 mM | 100 µM | No labeling (see below) | |

### SI5. Reaction between CSA (6a) and probes 3 and 5

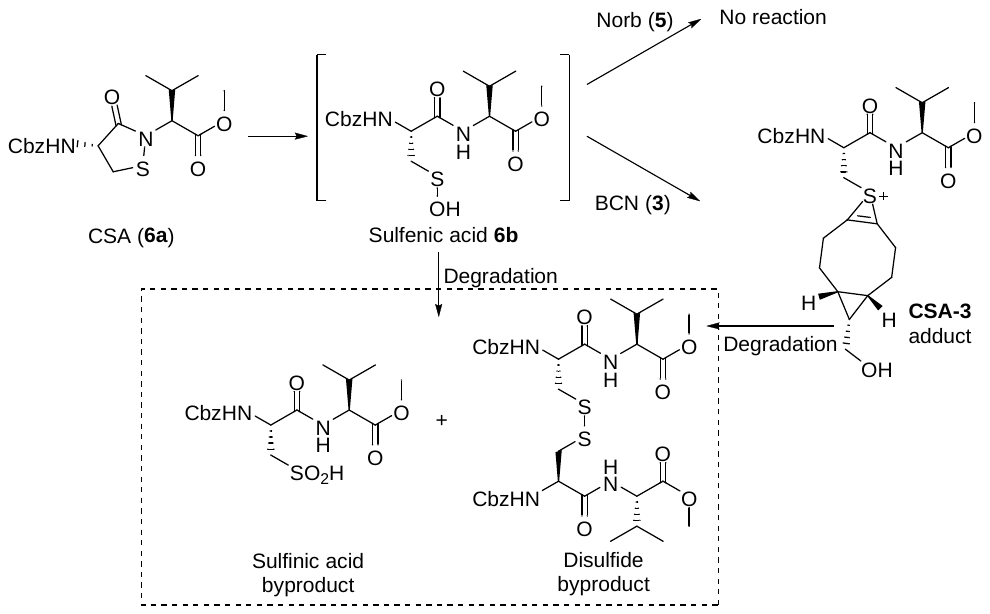

The reactions of probes BCN (**3**) and Norb (**5**) with CSA (**6a**) were performed following the procedures described above; however, these two probes did not yield stable adducts after 30 min.

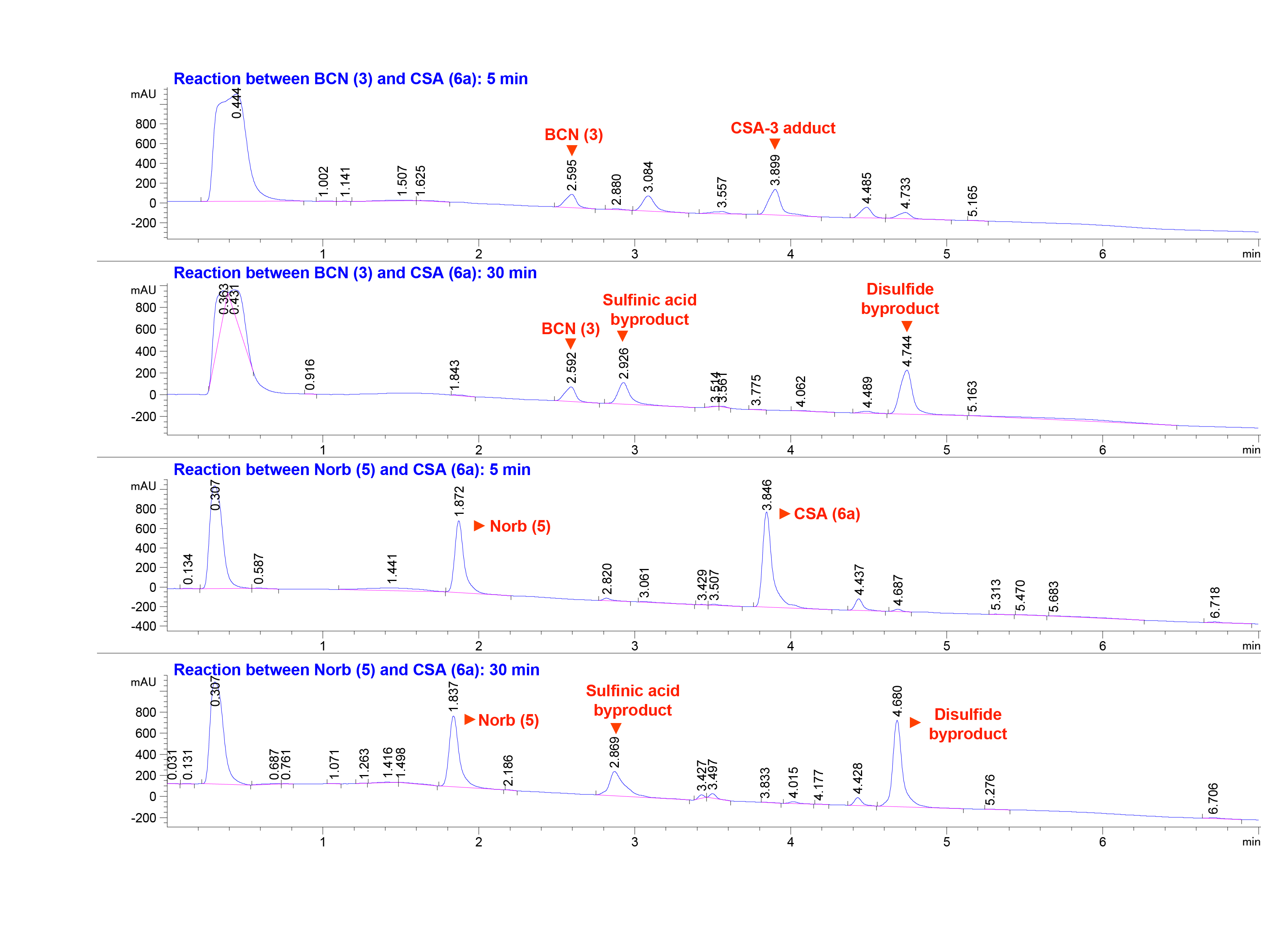

Figure S2. LC traces (190 nm) of probes **3** and **5** reacting with CSA (**6a**).

### SI6. Decaging of DMDE (7) and sulfenic acid capture with probes 1-5

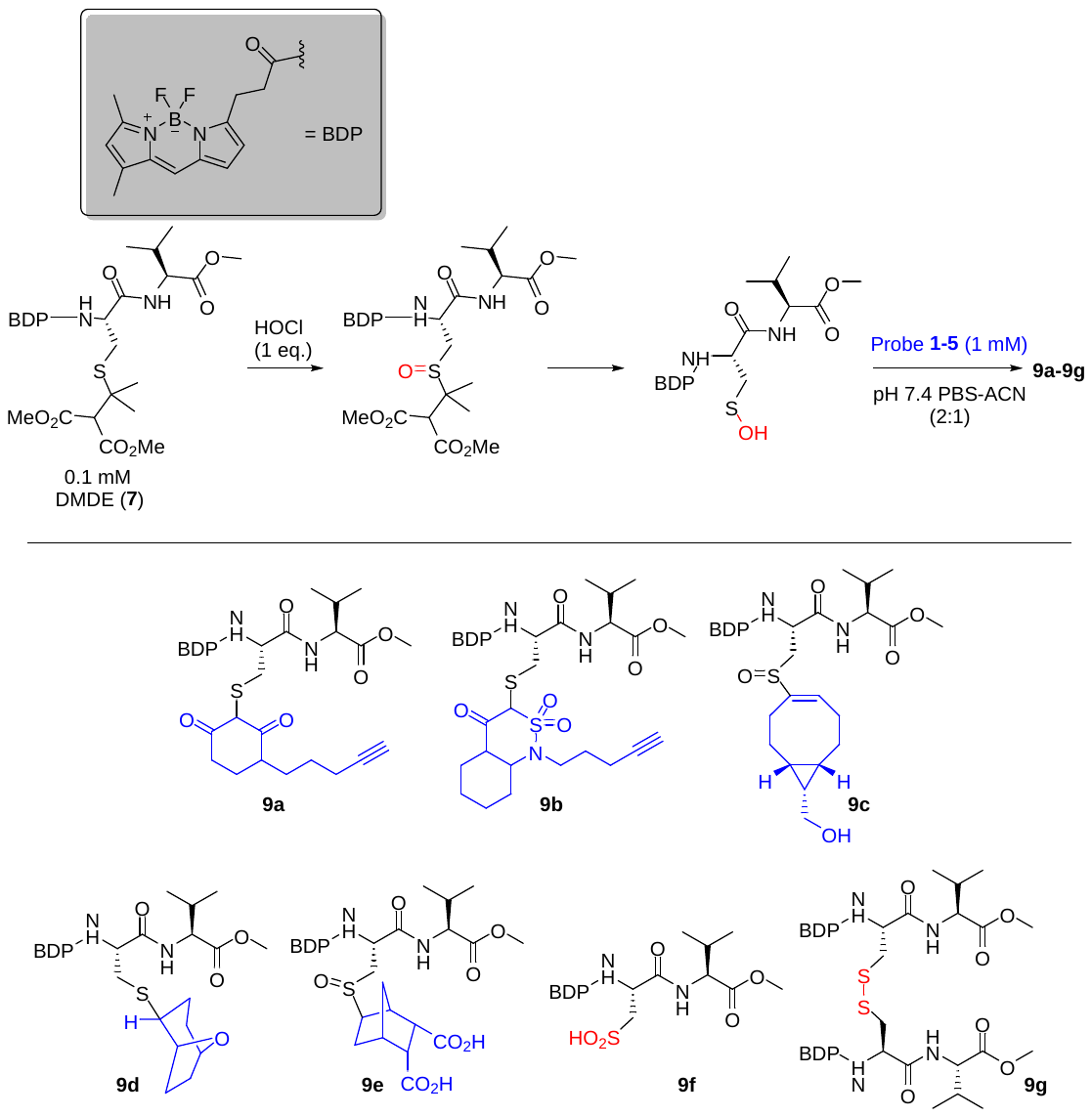

In pH 7.4 PBS-ACN buffer (2:1 v/v), 0.1 mM of DMDE (**7**) was first oxidized with 1 equivalent of HOCl to give the sulfoxide product, followed by addition of probes **1**-**5** (1 mM final concentration). After rocking at 37 °C for 16 h, the reactions were analyzed by LC-MS (493 nm absorption). Yield of each product **9a-g** was determined by integrated LC peak areas (using compound **7** as the standard), assuming all BODIPY-containing molecules have the same molar extinction coefficient.

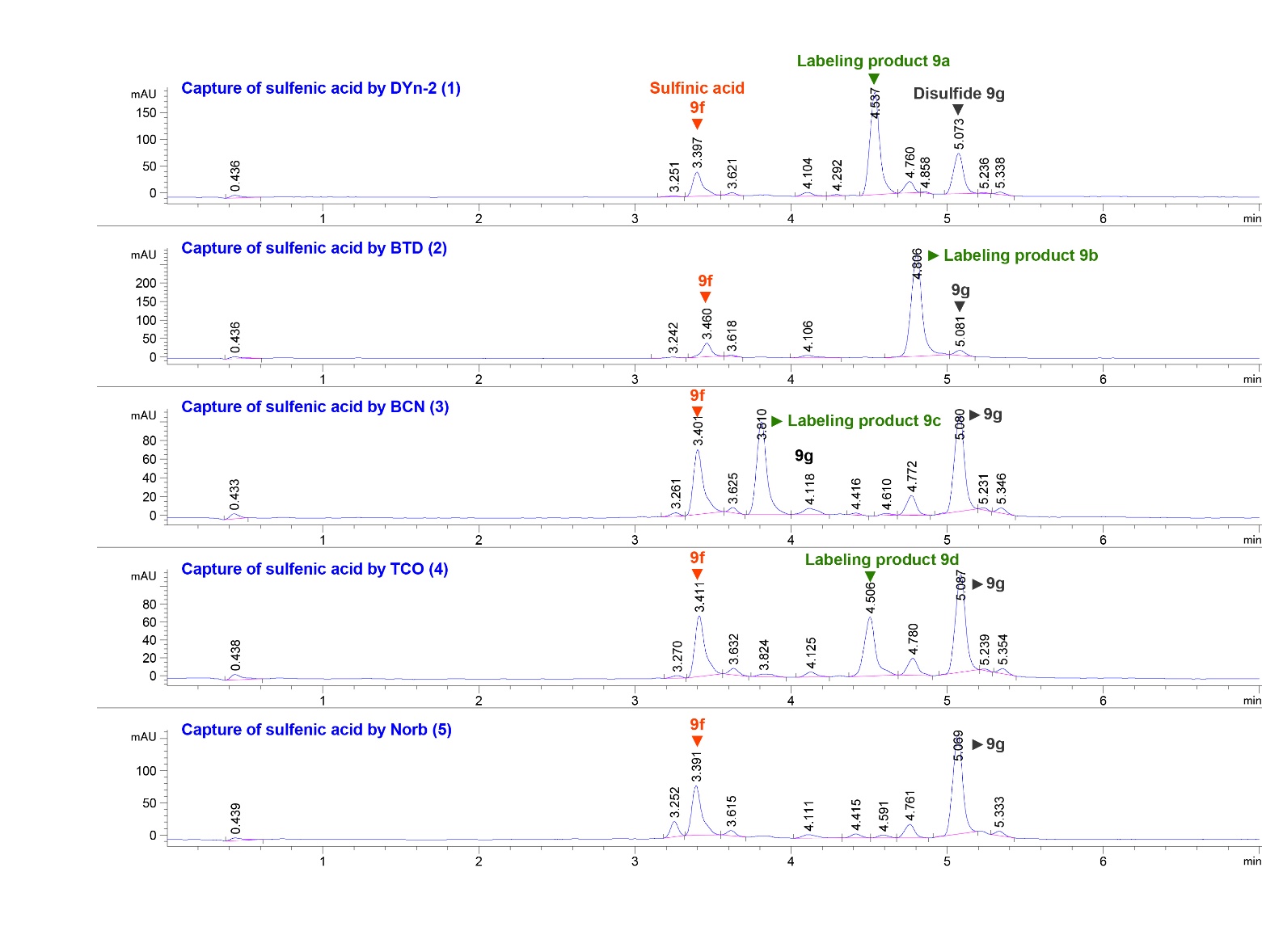

Figure S3. Representative LC-chromatograms (493 nm) of sulfenic acid capture from the fluorescent precursor **7** with probes **1-5**.

Table S2. Integrated peak areas that were used for calculating reaction yields.

| Peak area (mAU, 493 nm) | Labeling product 9a-9e | | Sulfinic acid 9f | | Disulfide 9g | |
| --- | --- | --- | --- | --- | --- | --- |
| Probes | Exp. #1 | Exp. #2 | Exp. #1 | Exp. #2 | Exp. #1 | Exp. #2 |
| DYn-2 (1) | 807.3 | 840 | 191.7 | 203 | 309.8 | 331.4 |
| BTD (2) | 1374 | 1341.5 | 162 | 157.5 | 66.2 | 61.7 |
| BCN (3) | 454 | 461.1 | 311.1 | 316.5 | 457.3 | 459.1 |
| TCO (4) | 345.5 | 334.4 | 313 | 306.9 | 501.3 | 491.7 |
| Norb (5) | 0 | 0 | 336 | 339.3 | 658.3 | 676.3 |

### SI7. Intact MS analyses of protein Gpx3 sulfenic acid with probes 1-5

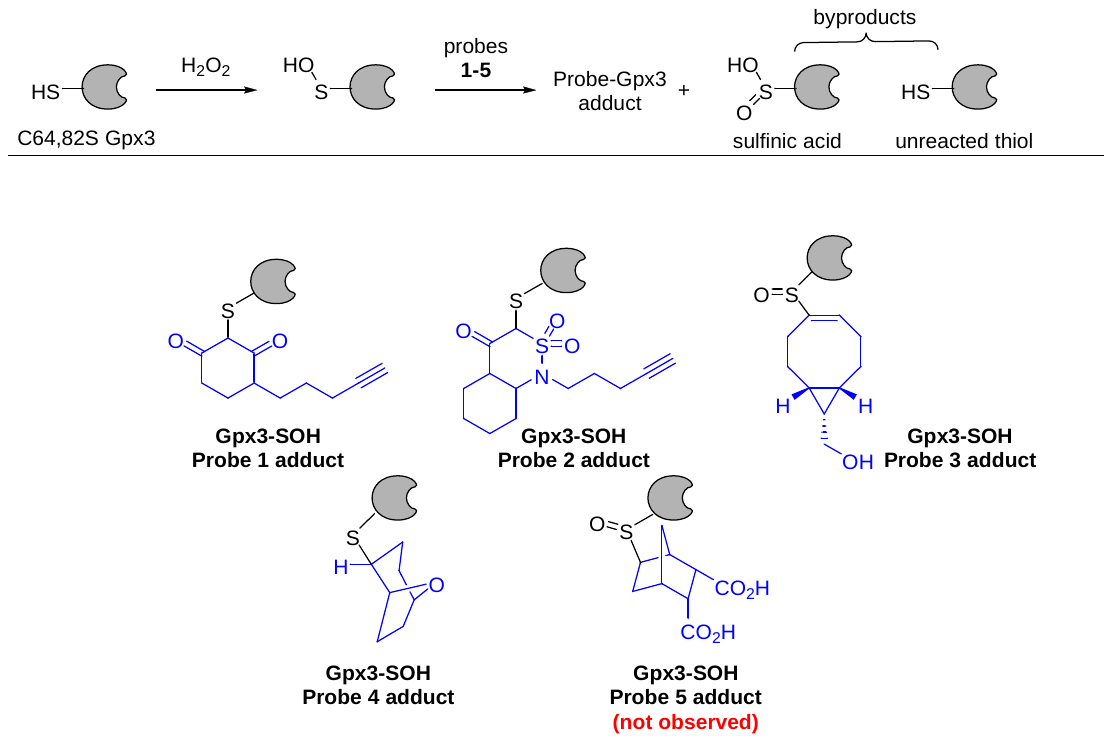

C64,82S Gpx3 stock solution (100 µM) was pre-reduced (50 mM DTT, 4 °C, 20 min) and desalted using a pre-equilibrated (HEPES buffer: 50 mM HEPES, 100 mM NaCl, pH 7.4) Nap-5 column (GE Healthcare). Protein concentration was determined using a Nanodrop (Ɛ = 24410 M^-1^·cm^-1^). The probes **1-5** (100 µM or 1 mM) was added to the diluted C64,82S Gpx3 solution (10 µM in HEPES buffer), which was subsequently treated with H_2_O_2_ (15 µM, 1.5 equiv.). The mixture (100 µl total volume) was incubated at room temperature for 1 h. After buffer exchange (50 mM ammonium bicarbonate), intact protein mass was acquired on a LTQ instrument (Thermo) after separation on a C4 protein column (50x4.6mm). Deconvoluted mass spectra were obtained using the MagTran software (V1.02, Z. Zhang, Amgen).

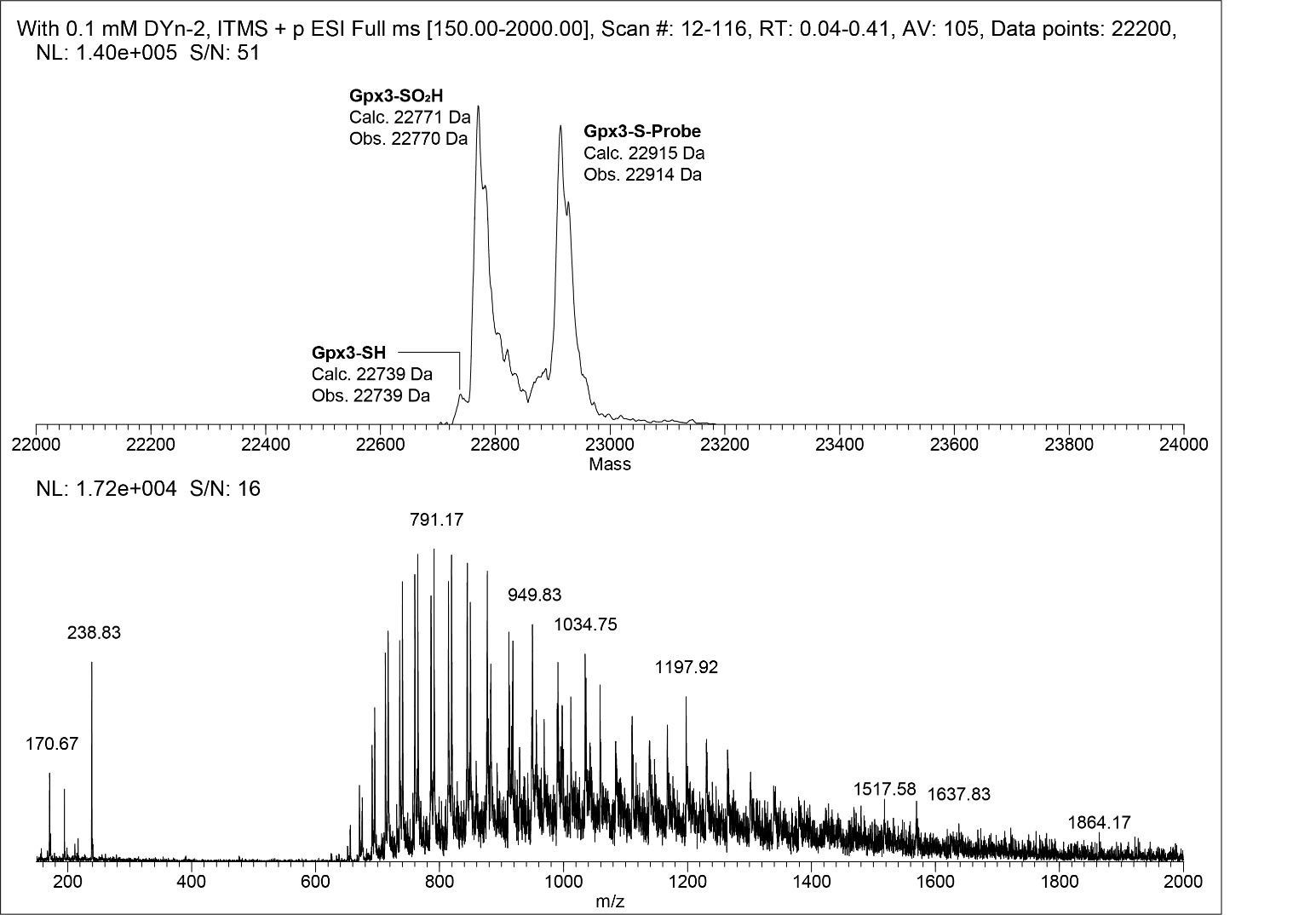

Figure S4. Raw and deconvoluted mass spectra of 10 µM Gpx3-SOH with 0.1 mM DYn-2.

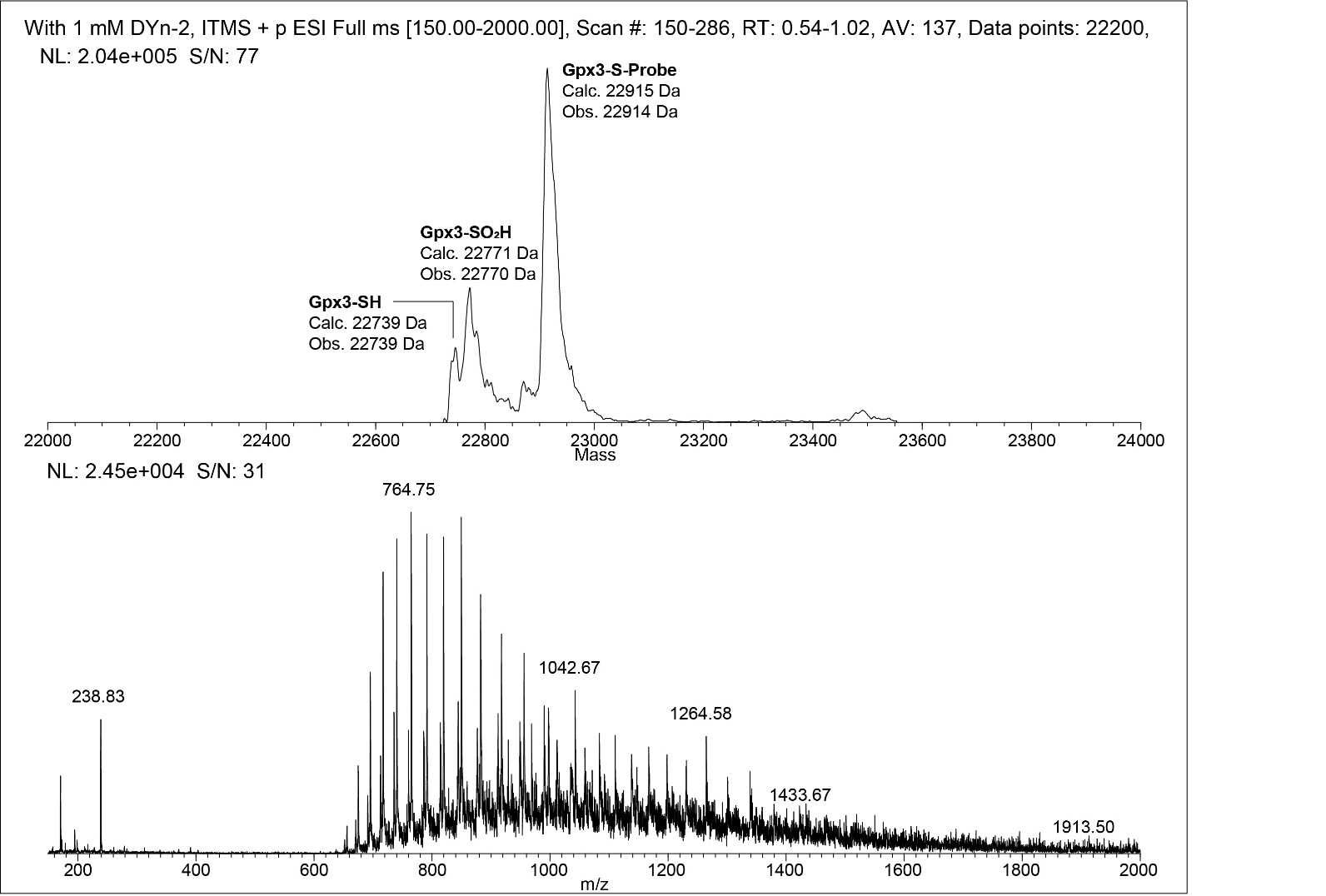

Figure S5. Raw and deconvoluted mass spectra of 10 µM Gpx3-SOH with 1 mM DYn-2.

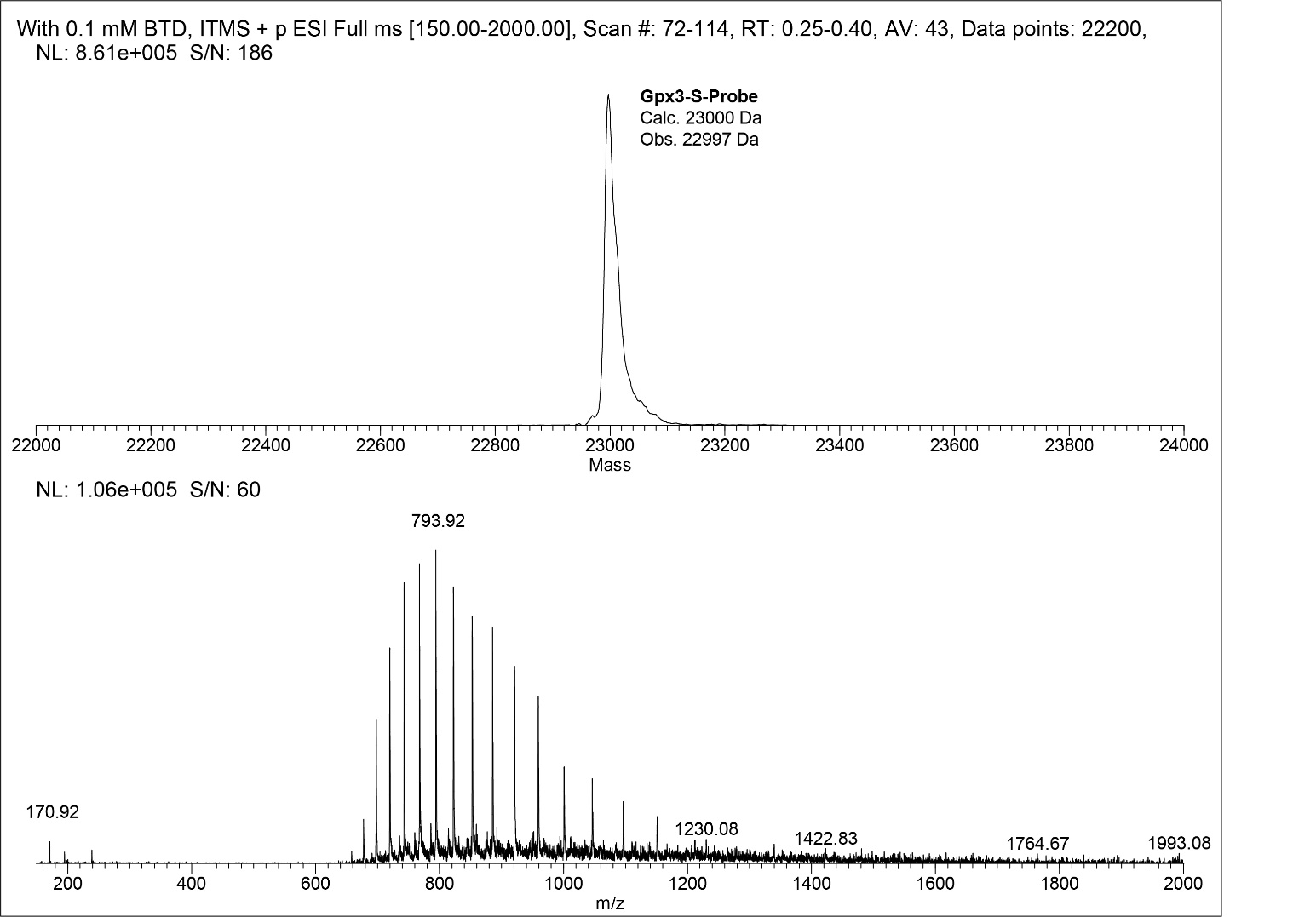

Figure S6. Raw and deconvoluted mass spectra of 10 µM Gpx3-SOH with 0.1 mM BTD.

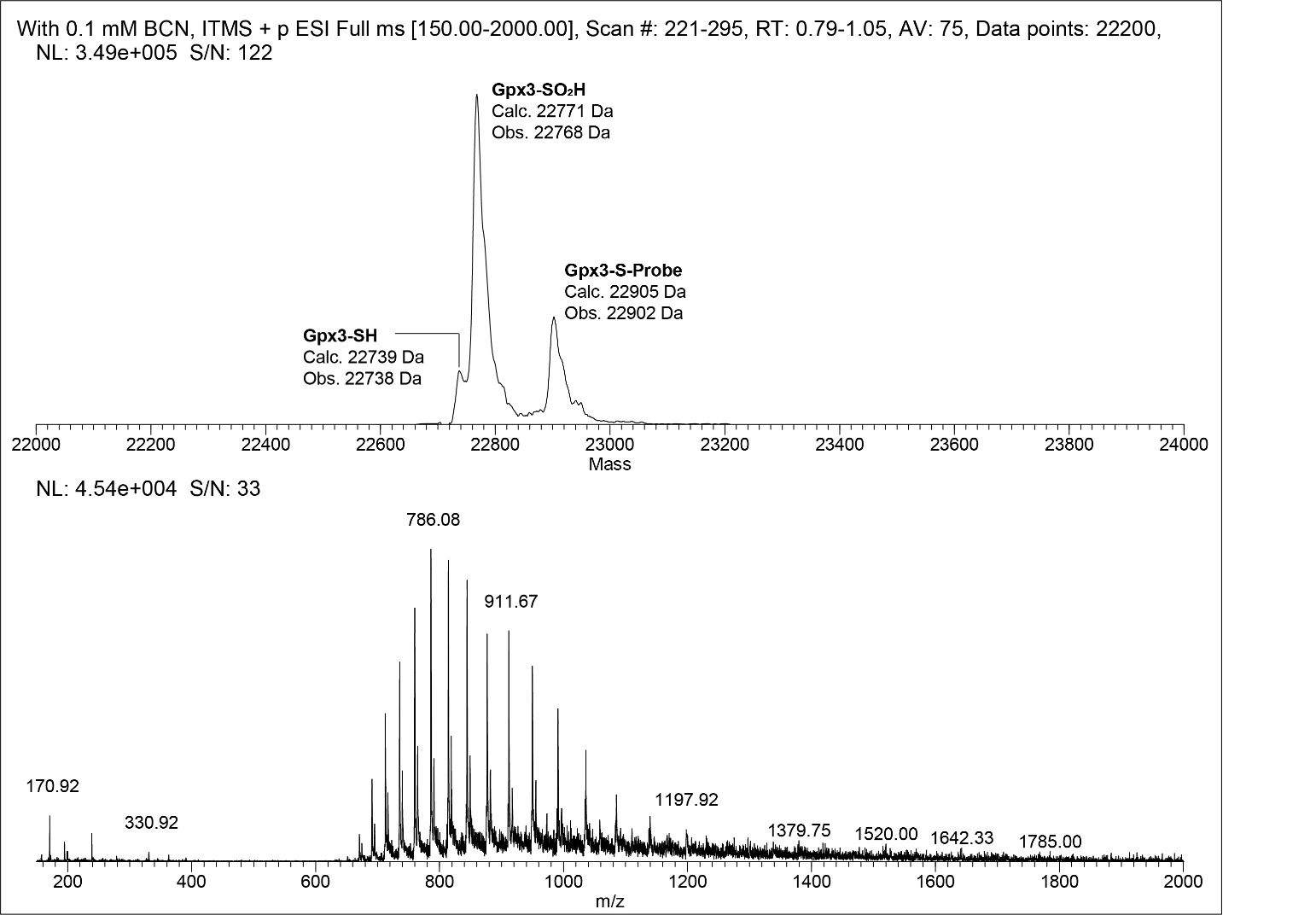

Figure S7. Raw and deconvoluted mass spectra of 10 µM Gpx3-SOH with 0.1 mM BCN.

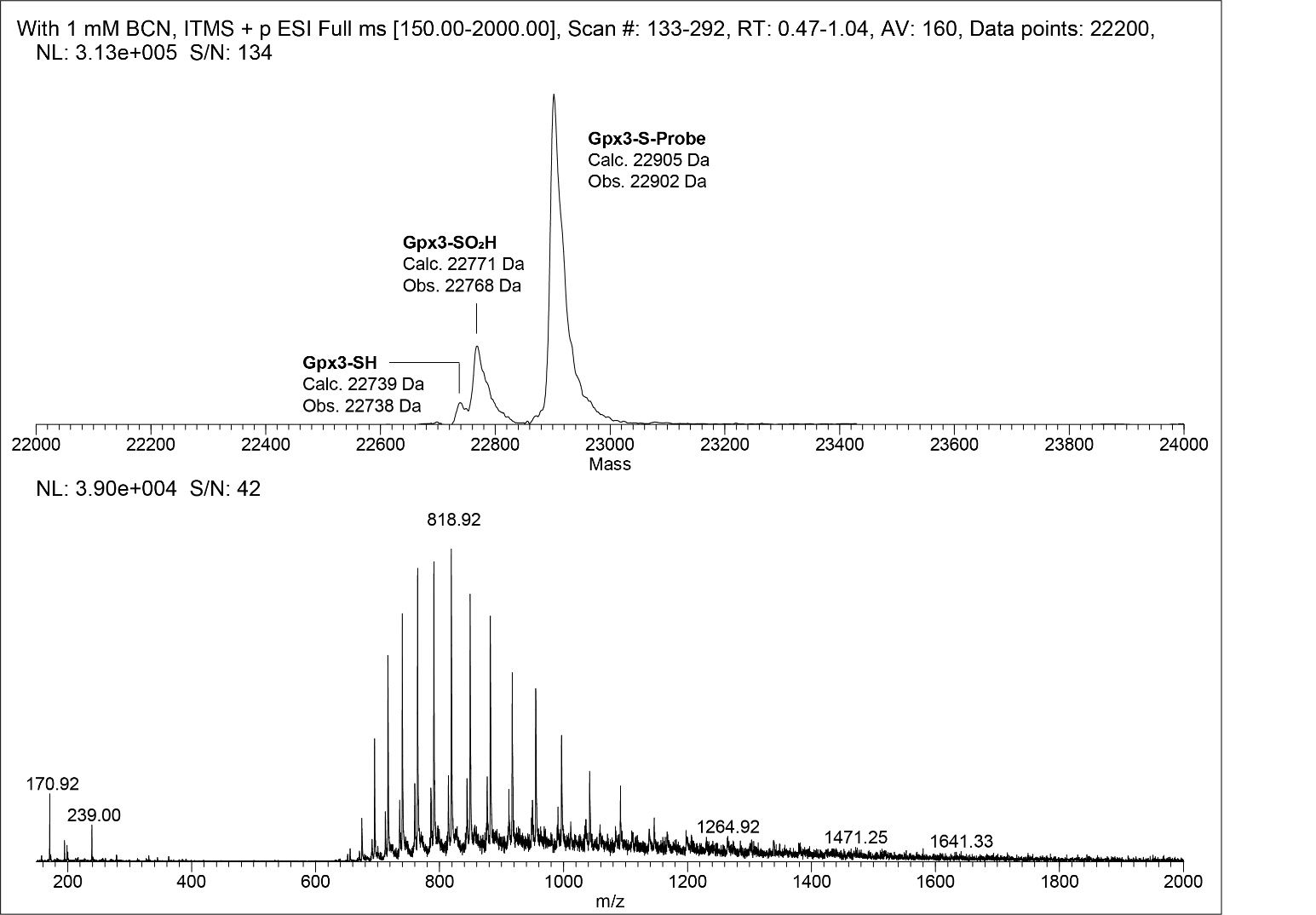

Figure S8. Raw and deconvoluted mass spectra of 10 µM Gpx3-SOH with 1 mM BCN.

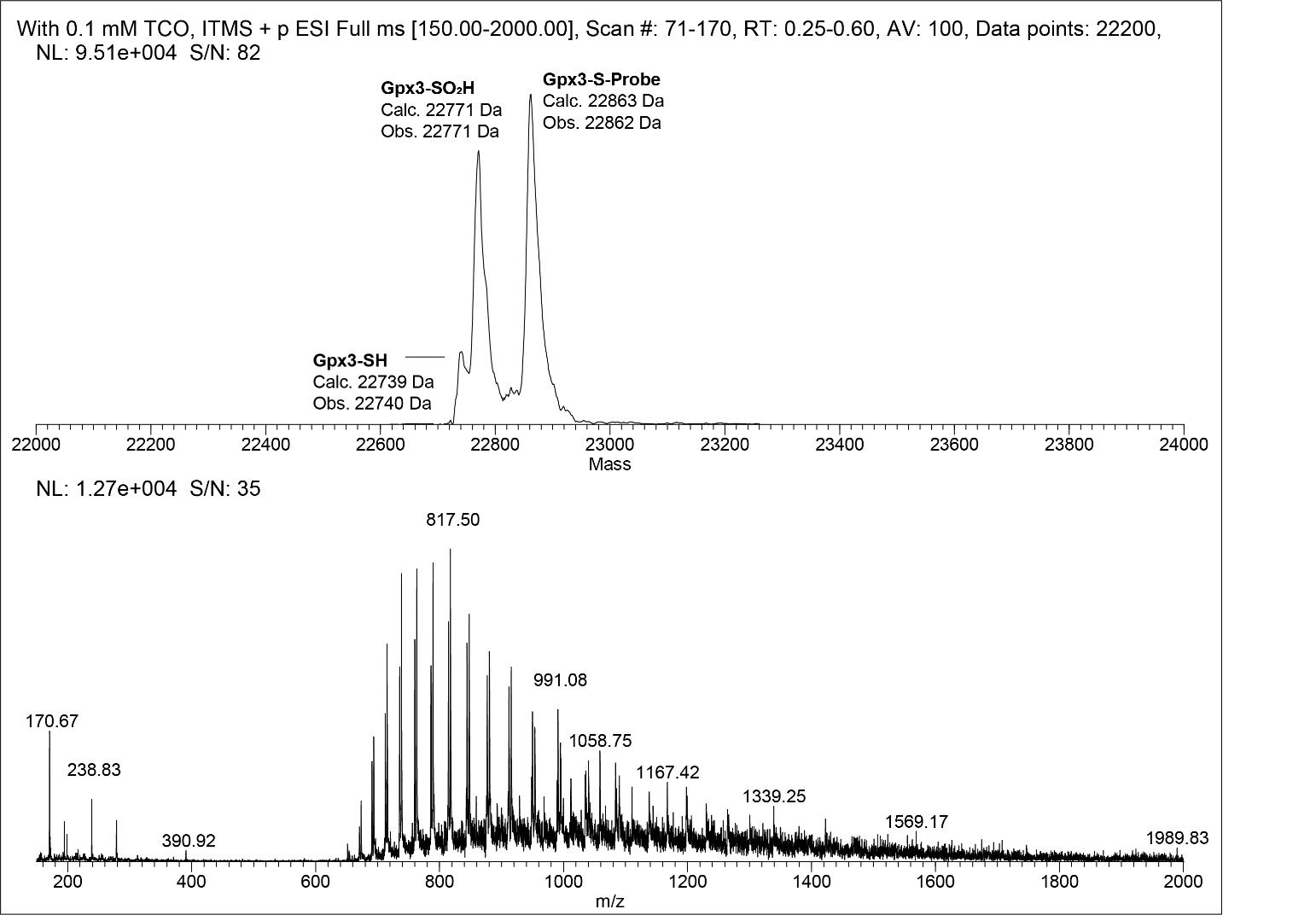

Figure S9. Raw and deconvoluted mass spectra of 10 µM Gpx3-SOH with 0.1 mM TCO.

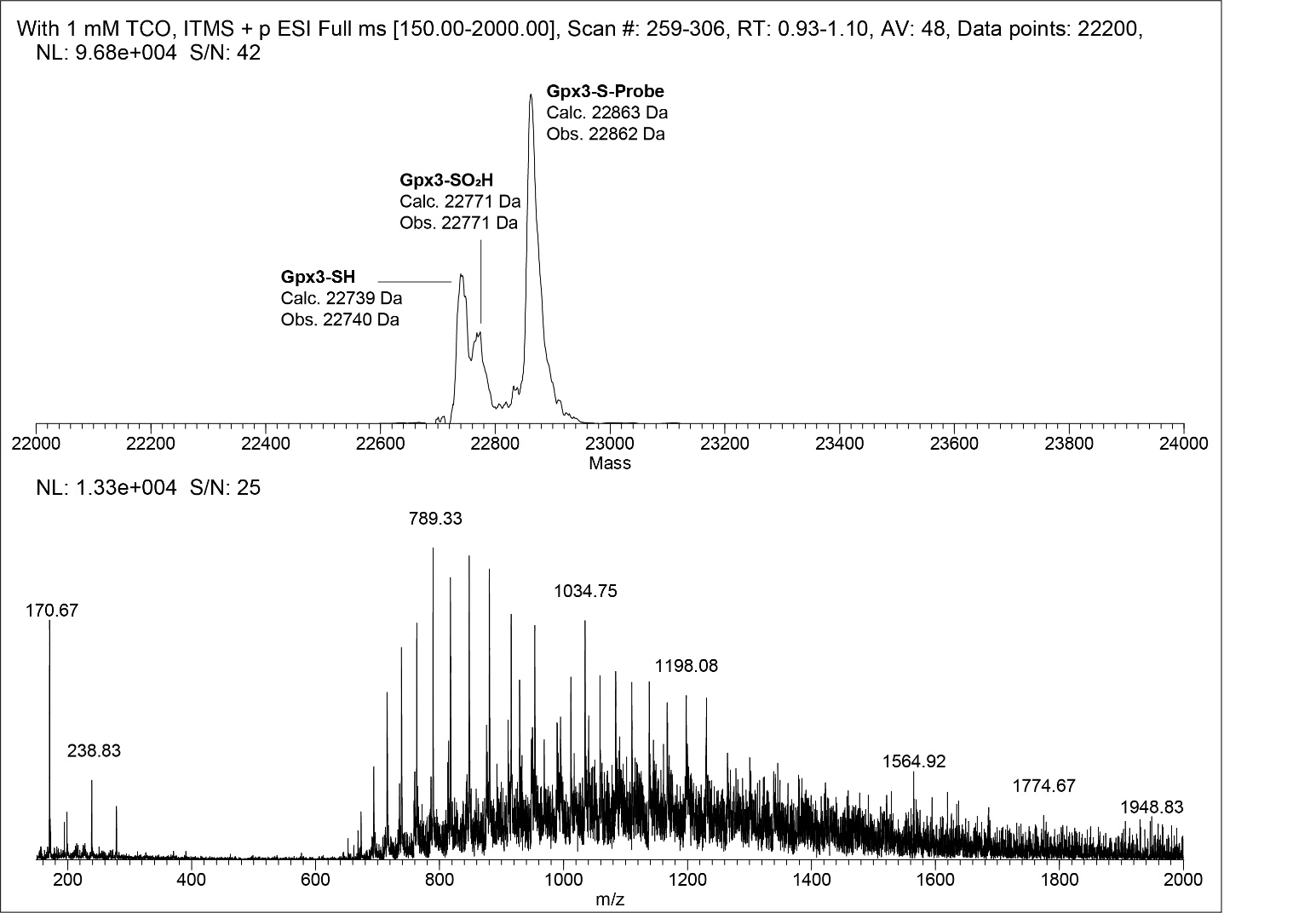

Figure S10. Raw and deconvoluted mass spectra of 10 µM Gpx3-SOH with 1 mM TCO.

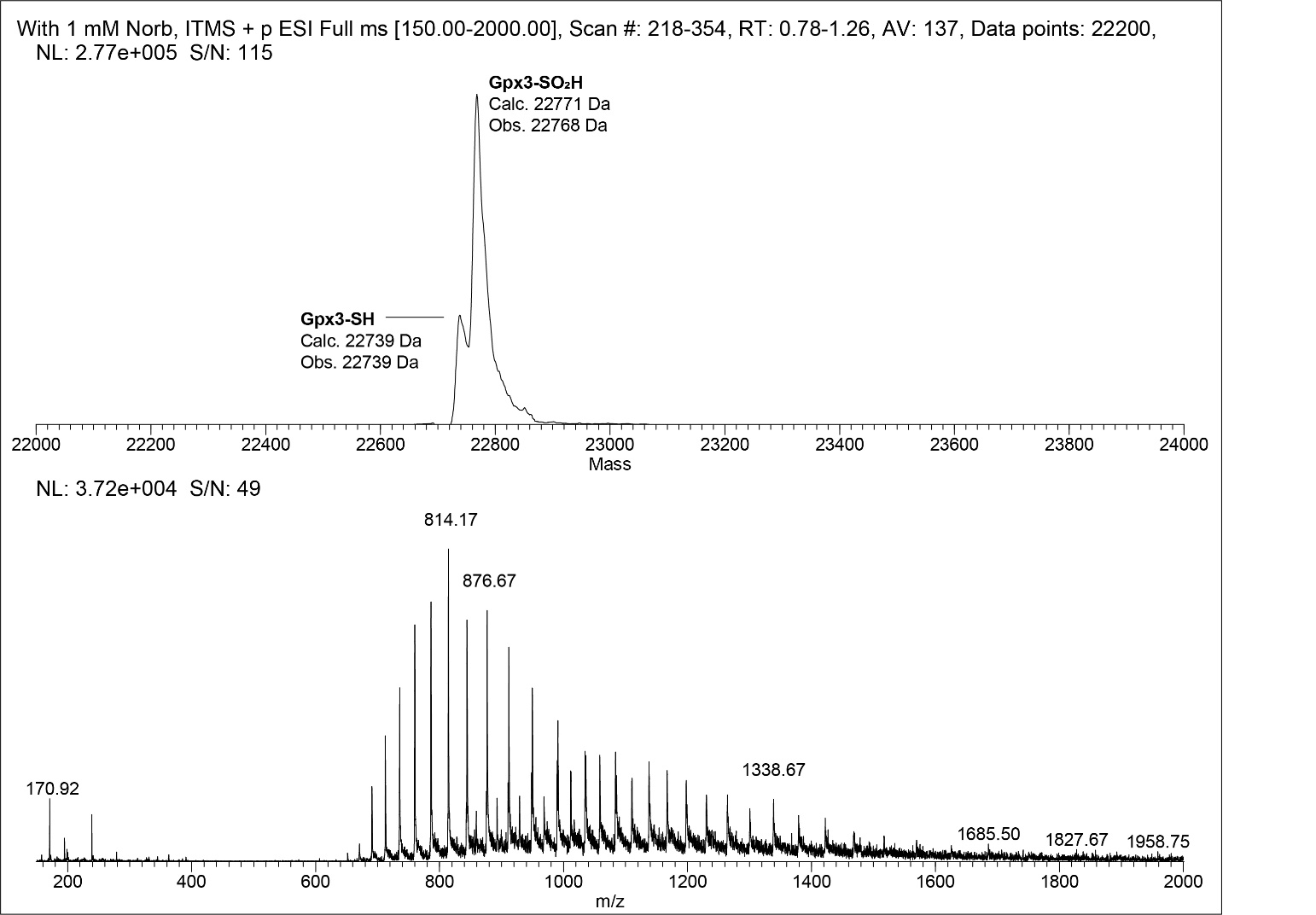

Figure S11. Raw and deconvoluted mass spectra of 10 µM Gpx3-SOH with 1 mM Norb.

### SI8. Intact MS analyses of oxidized protein PTP1B with nucleophiles

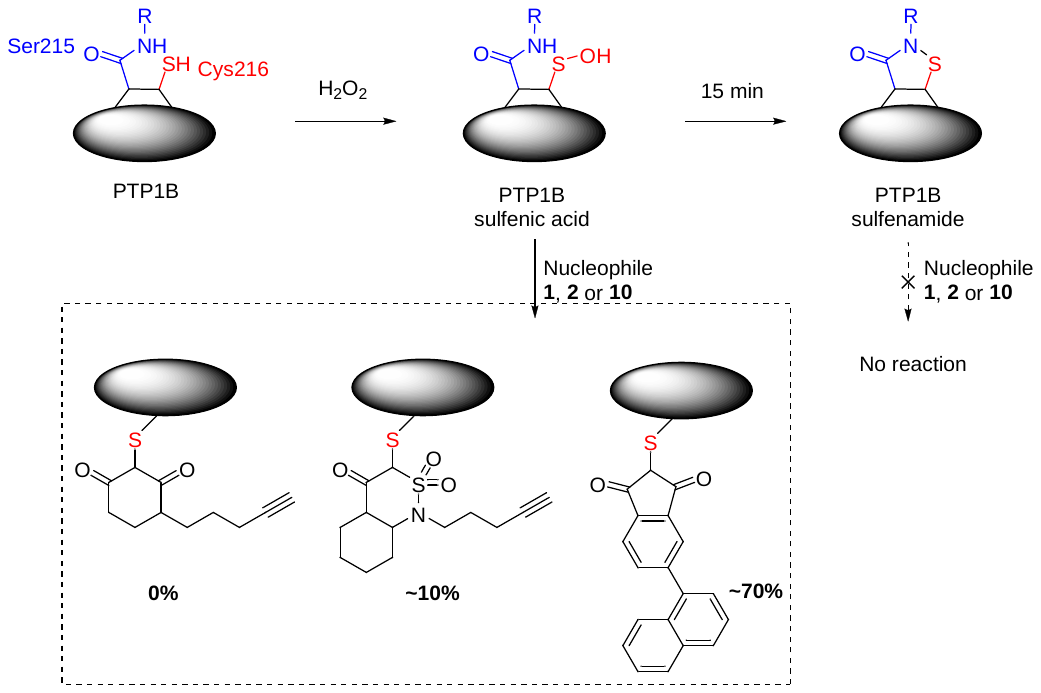

Stocks of PTP1B were treated with 50 mM DTT for 20 minutes on ice, followed by removal of DTT via buffer exchange with Nap-5 columns (GE Healthcare) that had been pre-equilibrated with labeling buffer (50 mM HEPES, 100 mM NaCl, 1 mM EDTA, pH 7.0). Protein concentrations were determined by A280 using a NanoDrop (Thermo). Protein samples were treated in one of two ways. In the first approach, we took 10 μM of PTP1B was treated with 1 mM of the nucleophile in the presence of 100 μM H_2_O_2_ for 1 hour. In the second approach, 10 μM PTP1B was pre-oxidized with 100 μM H_2_O_2_ for 15 minutes followed by the addition of 1 mM of the nucleophile. After buffer exchange (50 mM ammonium bicarbonate), intact protein mass was acquired on a LTQ instrument (Thermo) after separation on a C4 protein column (50x4.6mm). Deconvoluted mass spectra were obtained using the MagTran software (V1.02, Z. Zhang, Amgen).

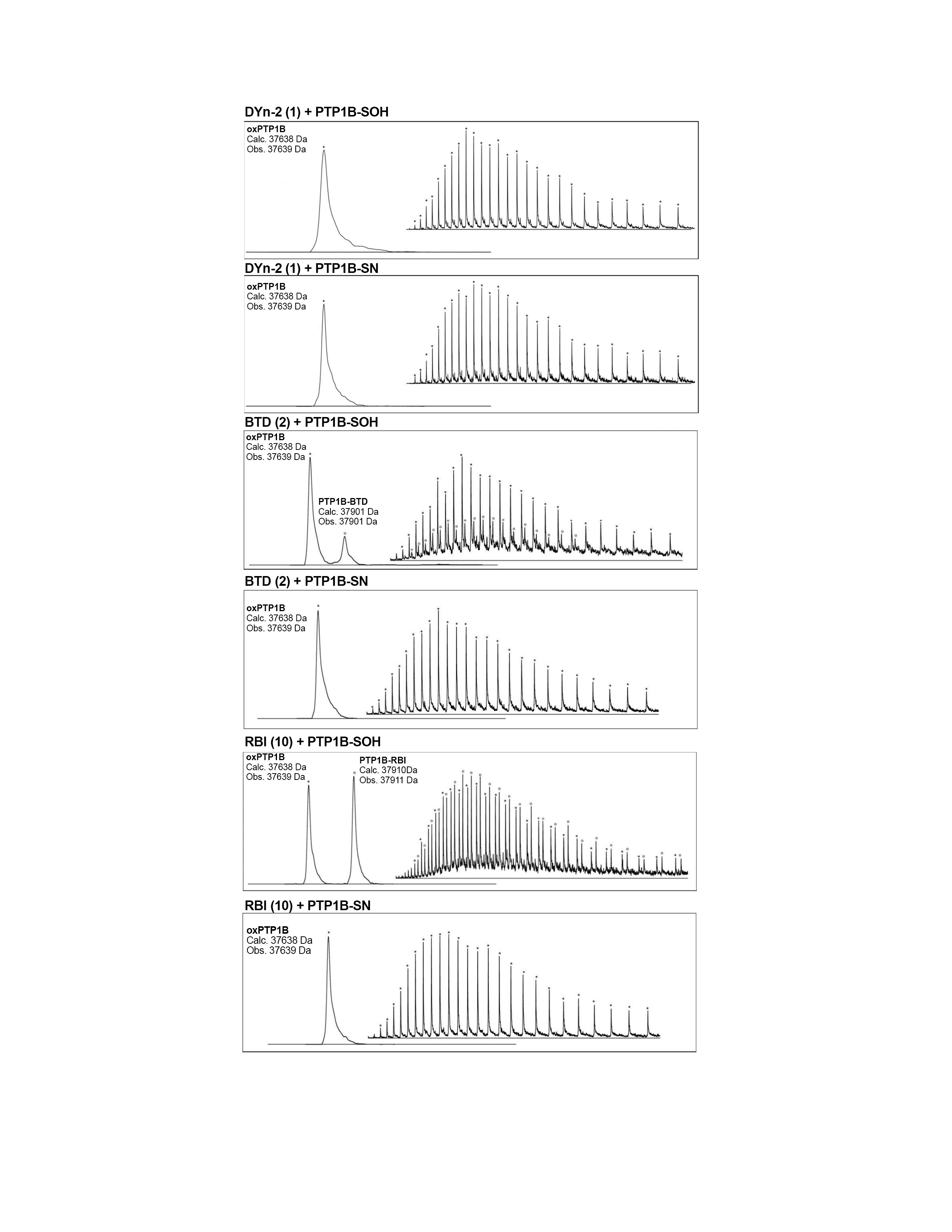

Figure S12. Raw and deconvoluted mass spectra of 10 µM PTP1B (pre-oxidized or oxidized in situ) with 1 mM nucleophiles **1**, **2** or **10**.

### SI9. Kinetic studies of sulfenamides with DYn-2 and BTD

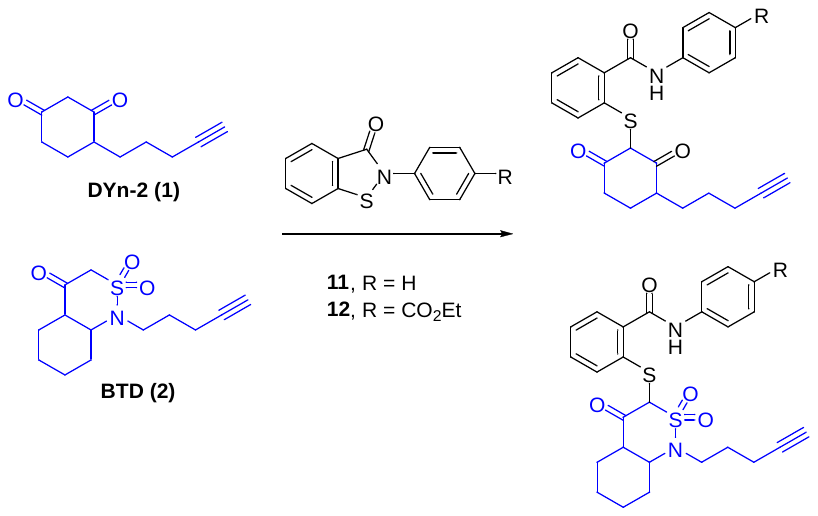

125 µM BTD or 1 mM DYn-2 (in 2:1 PBS(10 mM)-ACN pH 7.4) was treated with a benzisothiazolinone derivative **11** or **12** (25 µM for BTD; or 0.1 mM for DYn-2). Reaction was tracked at room temperature by LC-MS periodically for up to 20 h.

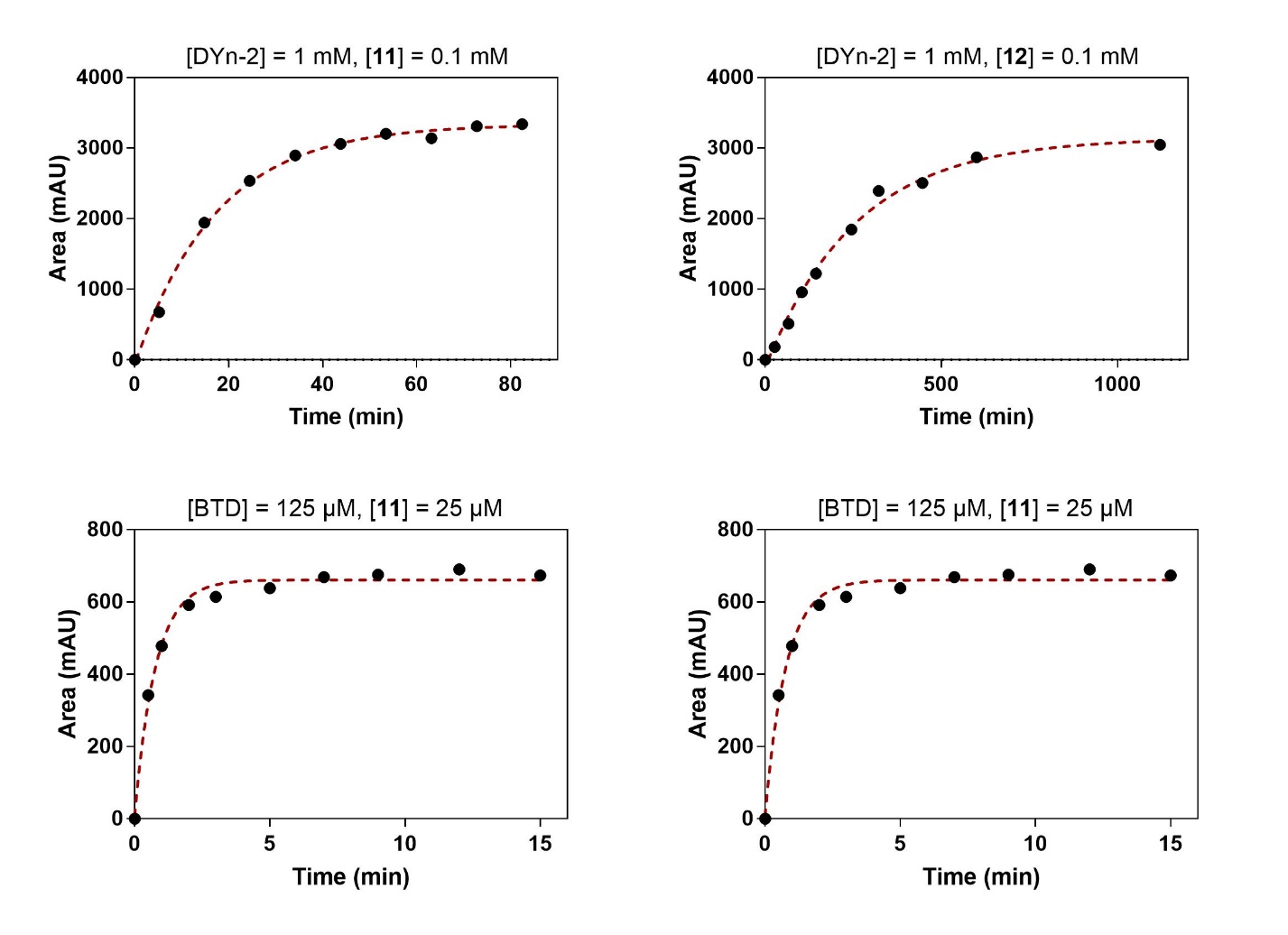

Figure S13. First order kinetic plots of probes **1** or **2** reacting with sulfenamides **11** or **12**.

Table S3. Calculations of probes **1** or **2** reacting with sulfenamides **11** or **12**.

| Probe | Probe conc. | Sulfenamide | Sulfenamide conc. | First order rate, *k* (min^-1^) | Calc’d 2^nd^ order rate, *K* (M^-1^·s^-1^) |
| --- | --- | --- | --- | --- | --- |
| DYn-2 (1) | 1 mM | **11** | 0.1 mM | 0.003873 | 0.06455 |
| DYn-2 (1) | 1 mM | **12** | 0.1 mM | 0.05777 | 0.9628 |
| BTD (2) | 125 µM | **11** | 25 µM | 0.08937 | 11.92 |
| BTD (2) | 125 µM | **12** | 25 µM | 1.288 | 171.7 |

### SI10. Kinetic studies of DYn-2 and BTD with various types of electrophiles

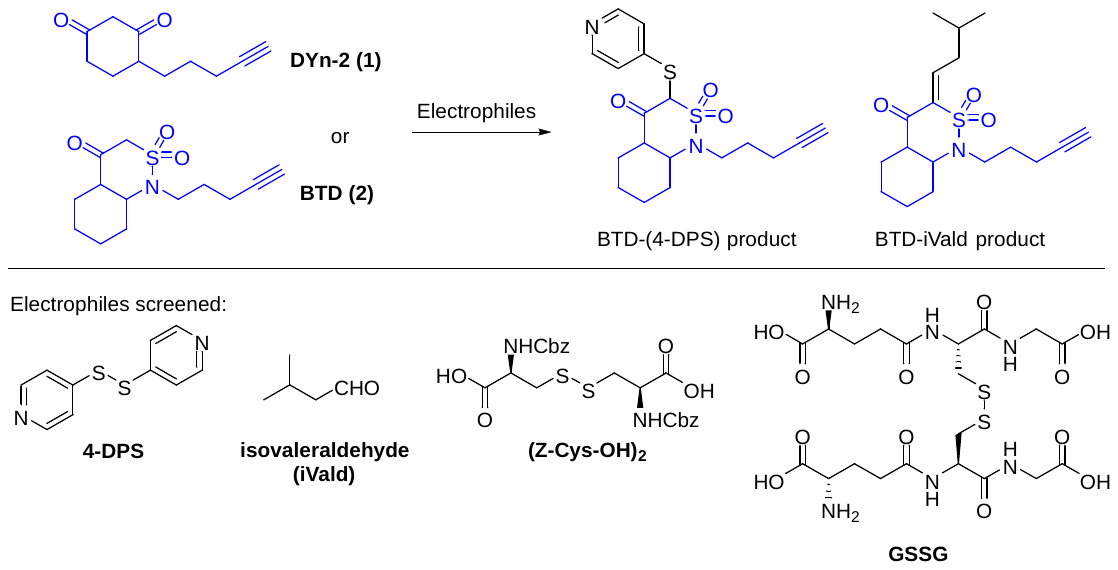

BTD or DYn-2 (5 mM in 2:1 PBS(10 mM)-ACN pH 7.4) was treated with 0.5 mM of electrophiles listed in the title (0.1 mM for isovaleraldehyde due to low solubility). Reaction was tracked at room temperature by LC-MS at 20 min interval for 120 min, then checked periodically for up to 24 h.

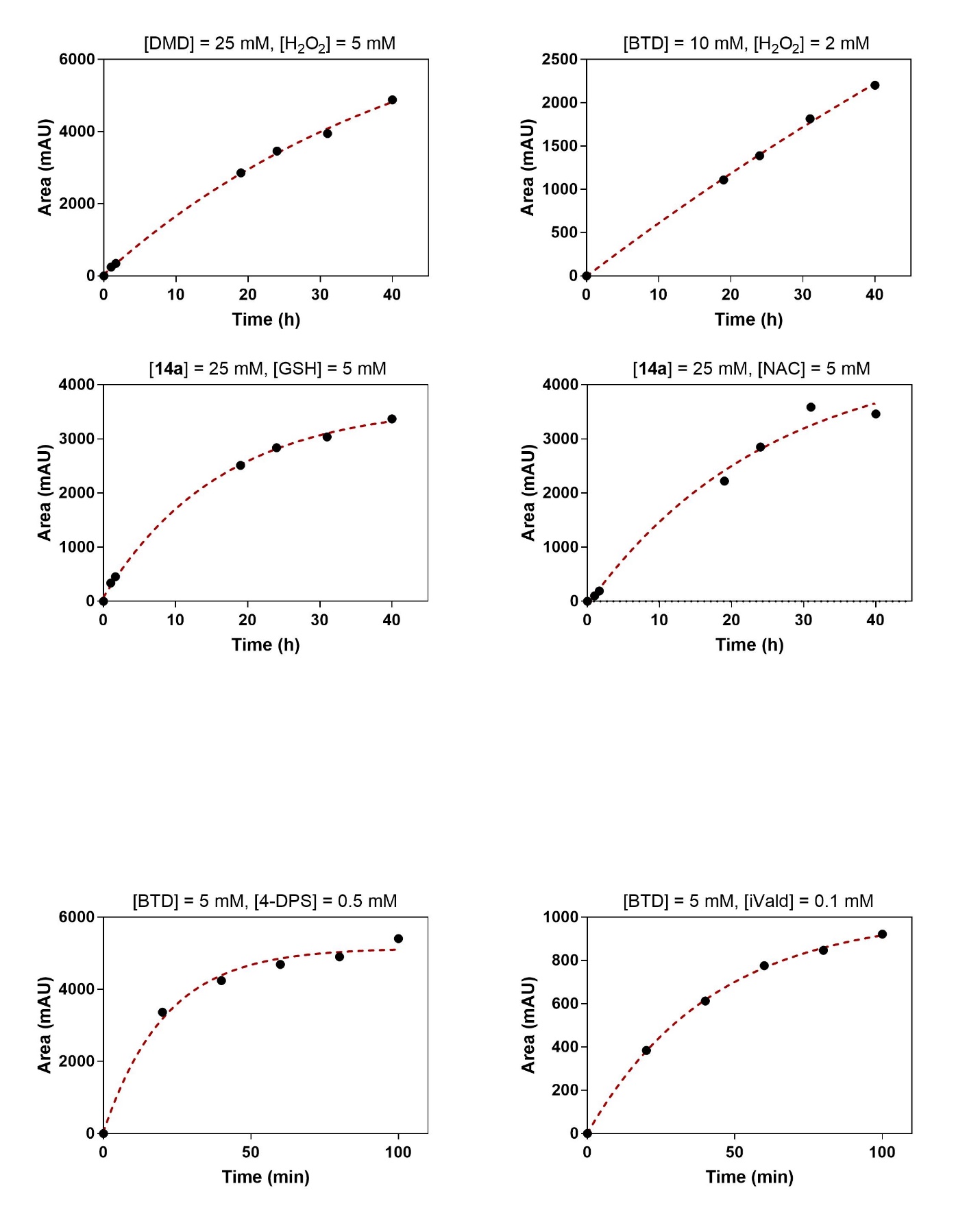

Figure S14. First order kinetic plots of **2** reacting with electrophiles 4-DPS or isovaleraldehyde.

Table S4. Calculations of **2** reacting with electrophiles.

| Probe | Probe conc. | Electrophile | Electrophile conc. | First order rate, *k* (min^-1^) | Calc’d 2^nd^ order rate, *K* (M^-1^·s^-1^) |
| --- | --- | --- | --- | --- | --- |
| DYn-2 (1) | 5 mM | 4-DPS | 0.5 mM | Incomplete (24 h) | < 1 x 10^-4^ |
| DYn-2 (1) | 5 mM | isovaleraldehyde | 0.1 mM | Incomplete (24 h) | < 1 x 10^-4^ |
| DYn-2 (1) | 5 mM | GSSG | 0.5 mM | No reaction | |
| DYn-2 (1) | 5 mM | (Z-Cys-OH)_2_ | 0.5 mM | No reaction | |
| BTD (2) | 5 mM | 4-DPS | 0.5 mM | 0.04777 | 0.1592 |
| BTD (2) | 5 mM | isovaleraldehyde | 0.1 mM | 0.02362 | 0.0787 |
| BTD (2) | 5 mM | GSSG | 0.5 mM | No reaction | |
| BTD (2) | 5 mM | (Z-Cys-OH)_2_ | 0.5 mM | No reaction | |

### SI11. Kinetic studies of probe oxidation and subsequent reactions with thiols

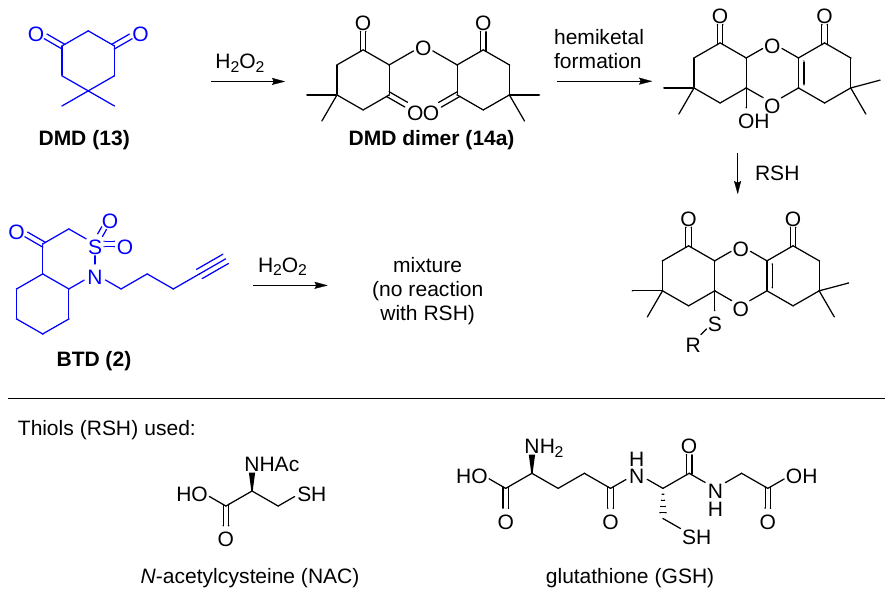

For the analysis of probe oxidation, 10 mM BTD or 25 mM dimedone (in 2:1 HEPES(50 mM)-ACN pH 7.4) was treated with hydrogen peroxide (2 mM or 5 mM, respectively). Reaction was tracked at room temperature by LC-MS periodically for up to 40 h.

For the reaction of DMD dimer with thiols, 25 mM of purified DMD dimer (**14a** in 2:1 HEPES(50 mM)-ACN pH 7.4) was treated with 5 mM of *N*-acetylcysteine (NAC) or glutathione (GSH). Reaction was tracked at room temperature by LC-MS periodically for up to 40 h.

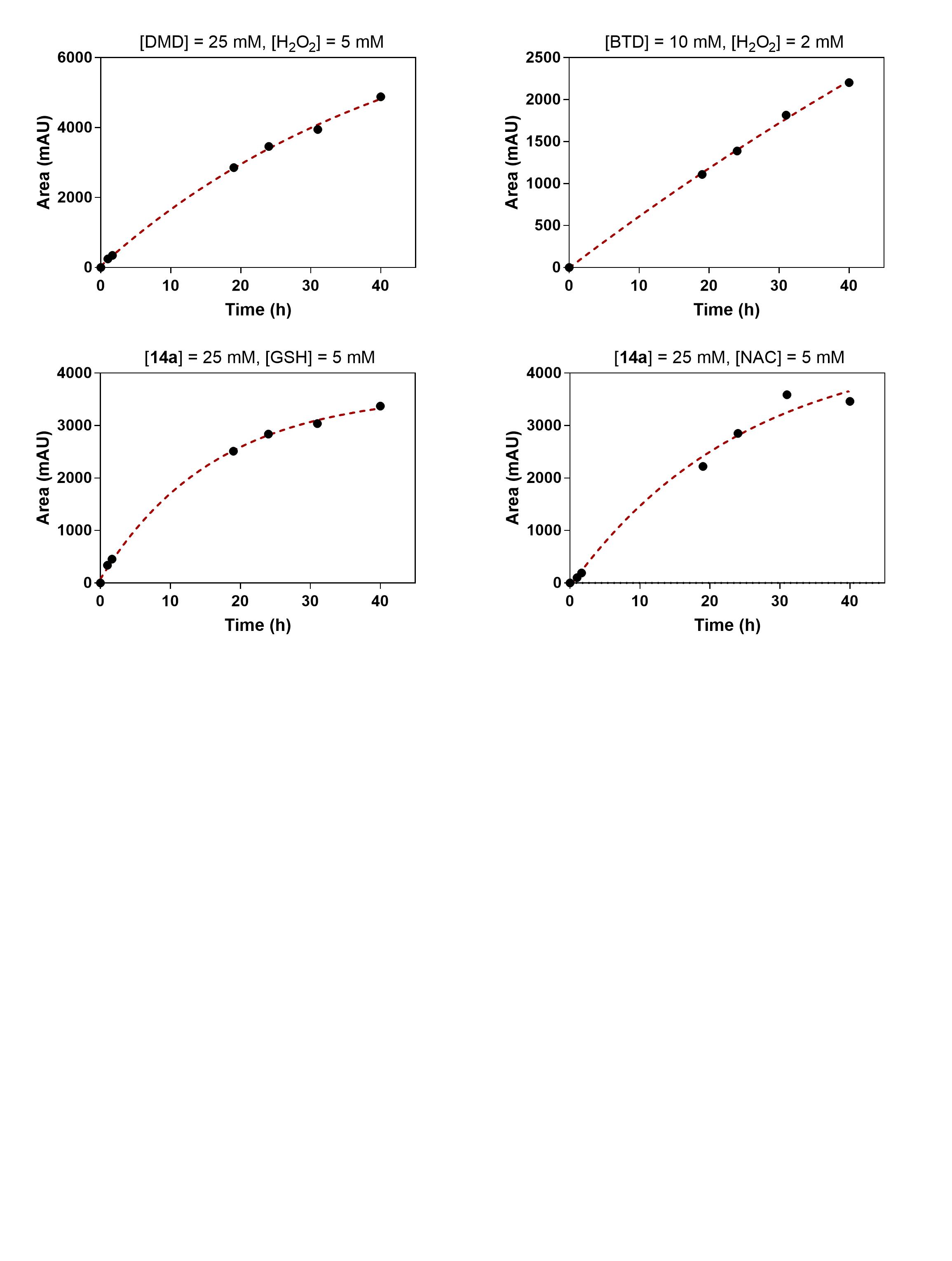

Figure S15. First order kinetic plots of probe oxidation and subsequent reactions with thiols.

Table S5. Calculations of probe oxidation and subsequent reactions with thiols.

| Substrate | Substrate conc. | Additive | Additive conc. | First order rate, *k* (h^-1^) | Calc’d 2^nd^ order rate, *K* (M^-1^·s^-1^) |
| --- | --- | --- | --- | --- | --- |
| DMD (13) | 25 mM | H_2_O_2_ | 5 mM | 0.02234 | 2.482 x 10^-4^ |
| BTD (2) | 10 mM | H_2_O_2_ | 2 mM | 0.00659 | 1.831 x 10^-4^ |
| DMD dimer (14a) | 25 mM | GSH | 5 mM | 0.06126 | 6.807 x 10^-4^ |
| DMD dimer (14a) | 25 mM | NAC | 5 mM | 0.0395 | 4.422 x 10^-4^ |

### NMR spectra

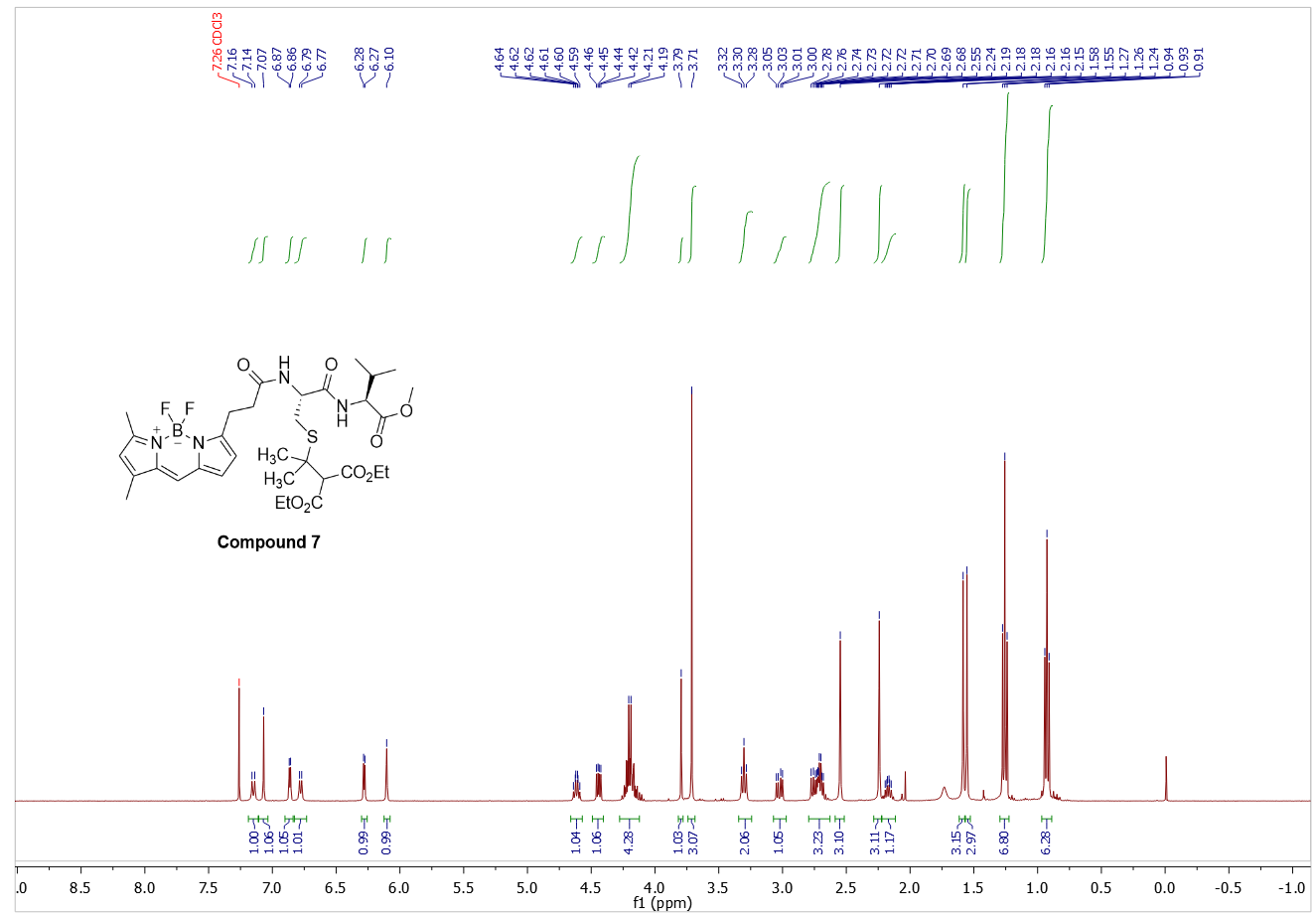
**^1^H NMR of compound 7**

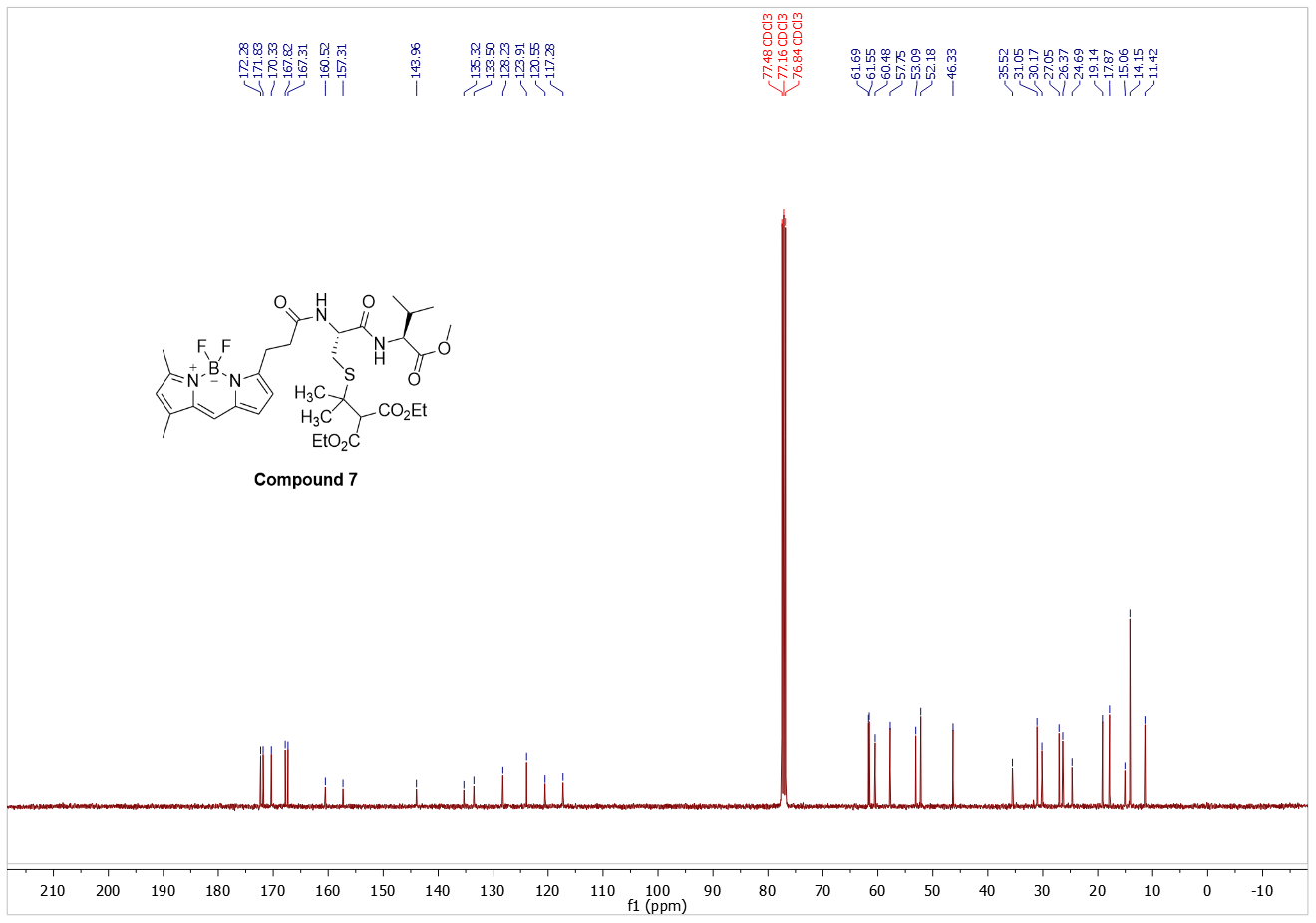

**^13^C NMR of compound 7**

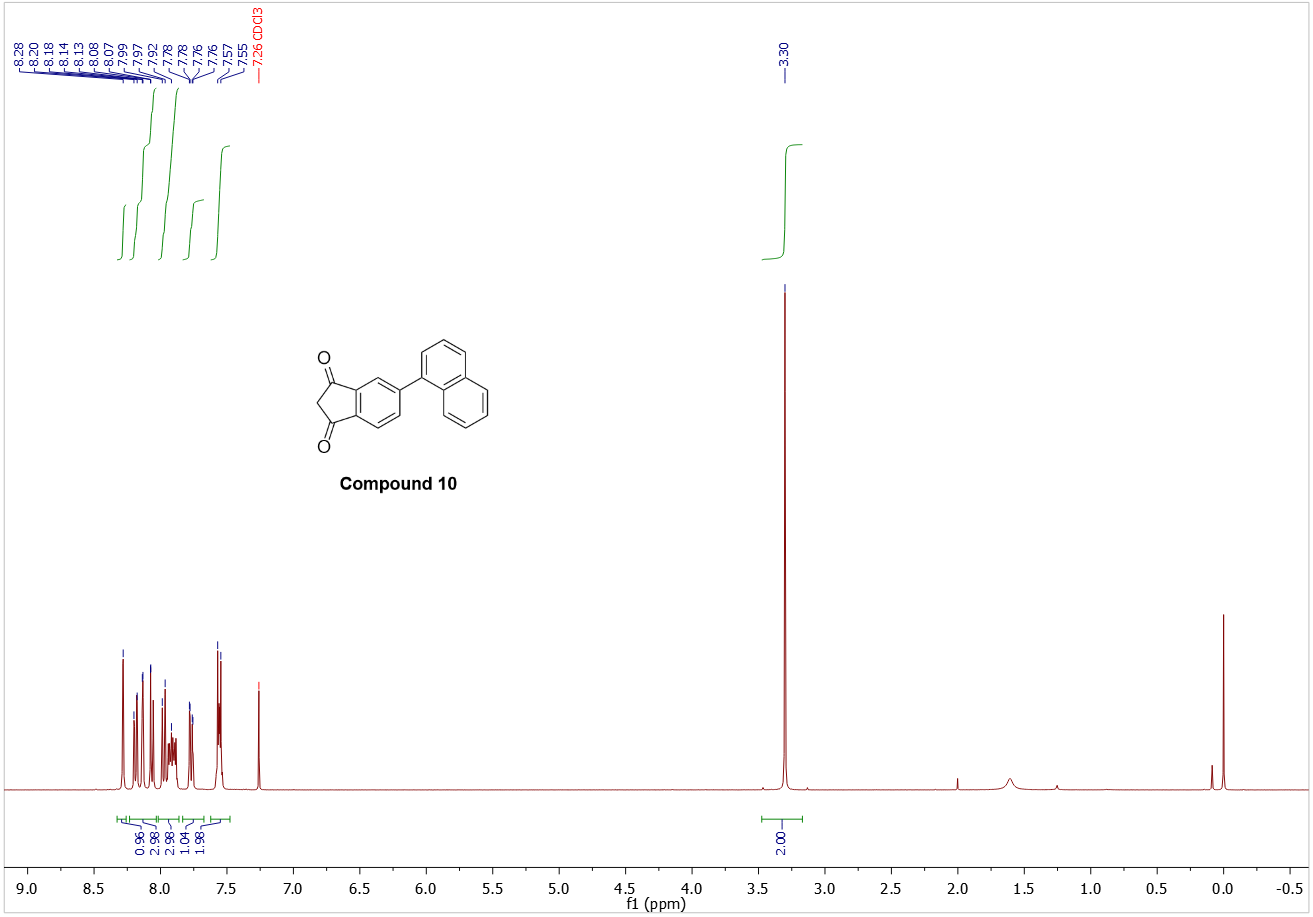

**^1^H NMR of compound 10**

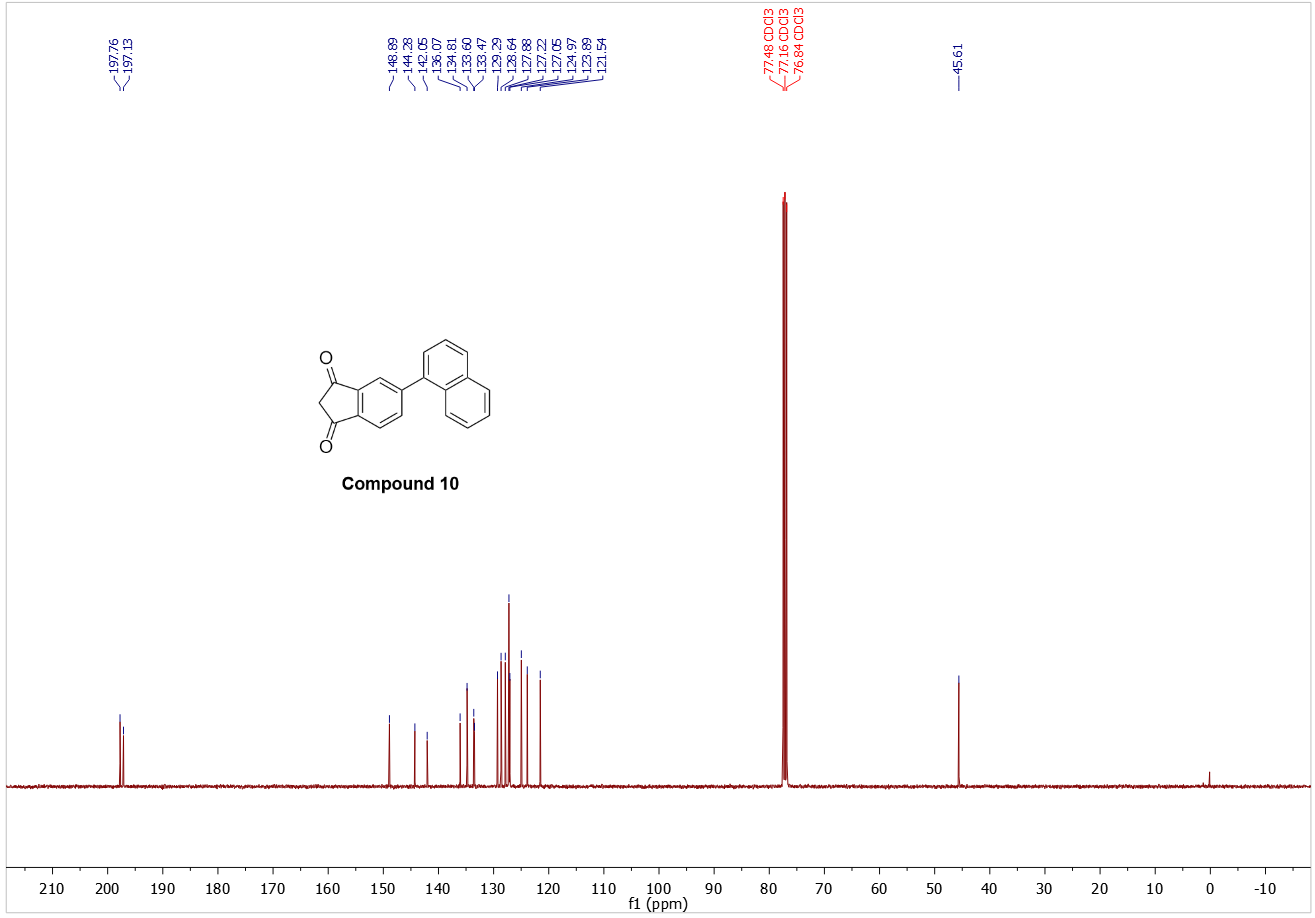

**^13^C NMR of compound 10**

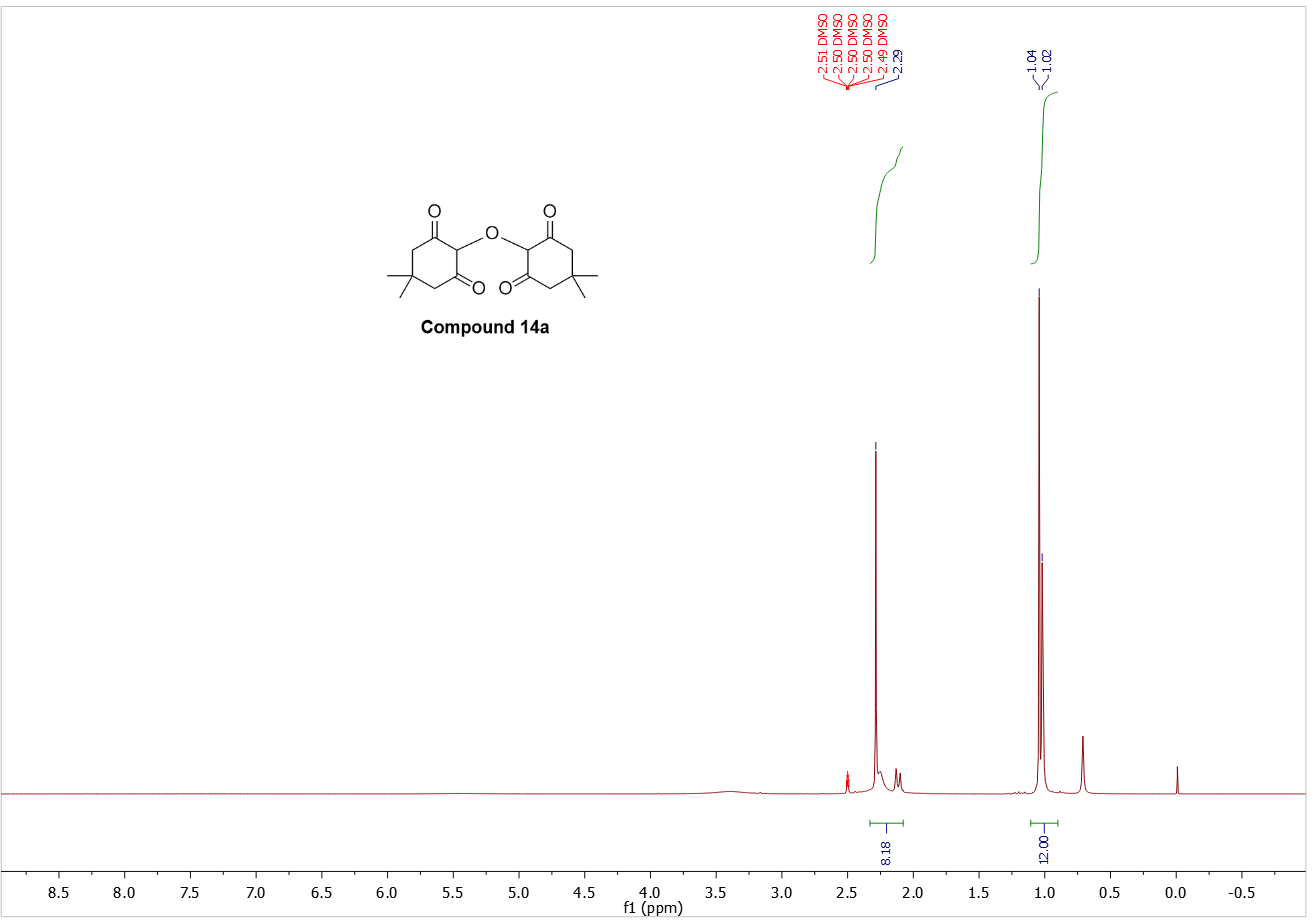

**^1^H NMR of compound 14a (in DMSO-d_6_)**

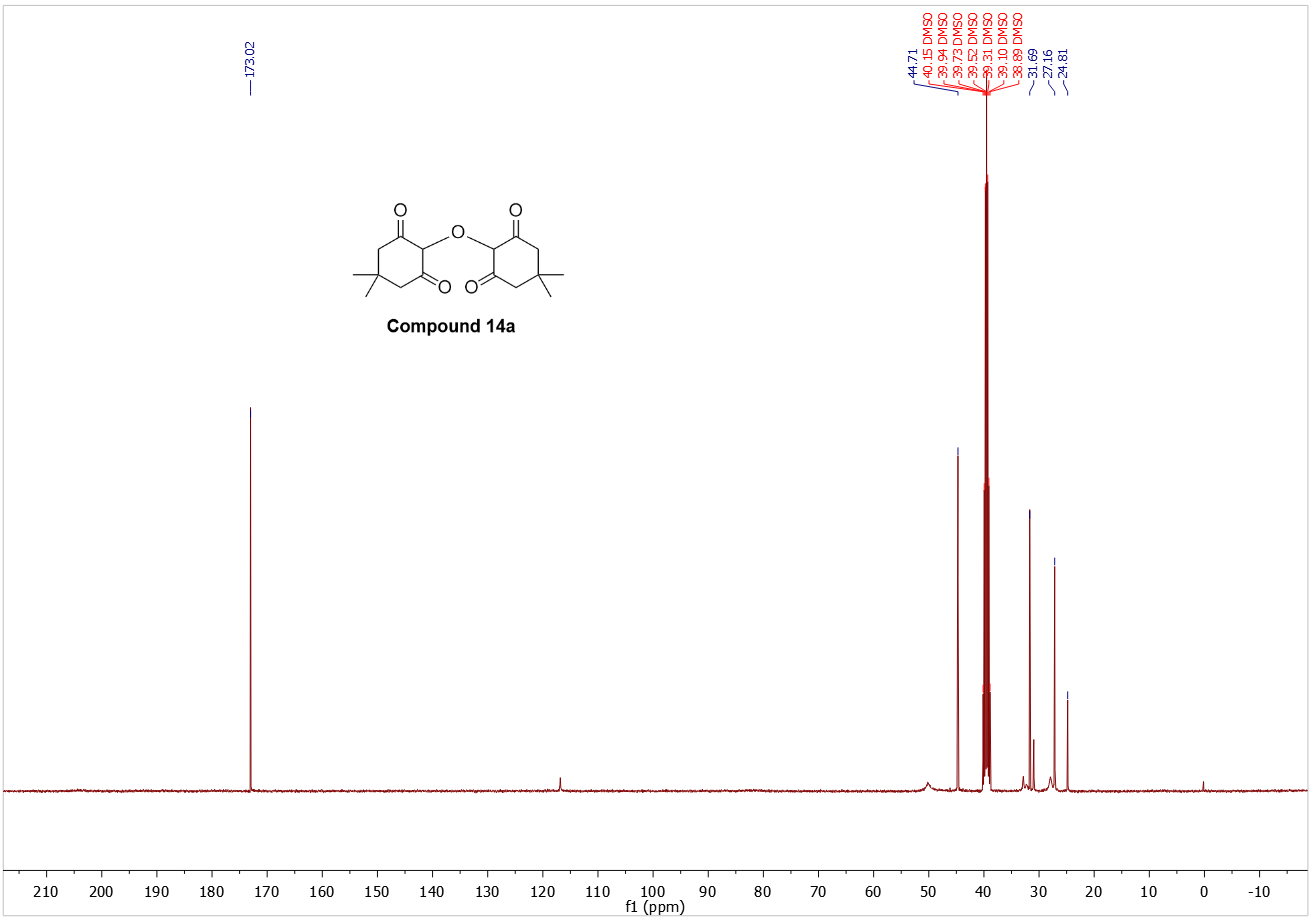

**^13^C NMR of compound 14a (in DMSO-d_6_)**

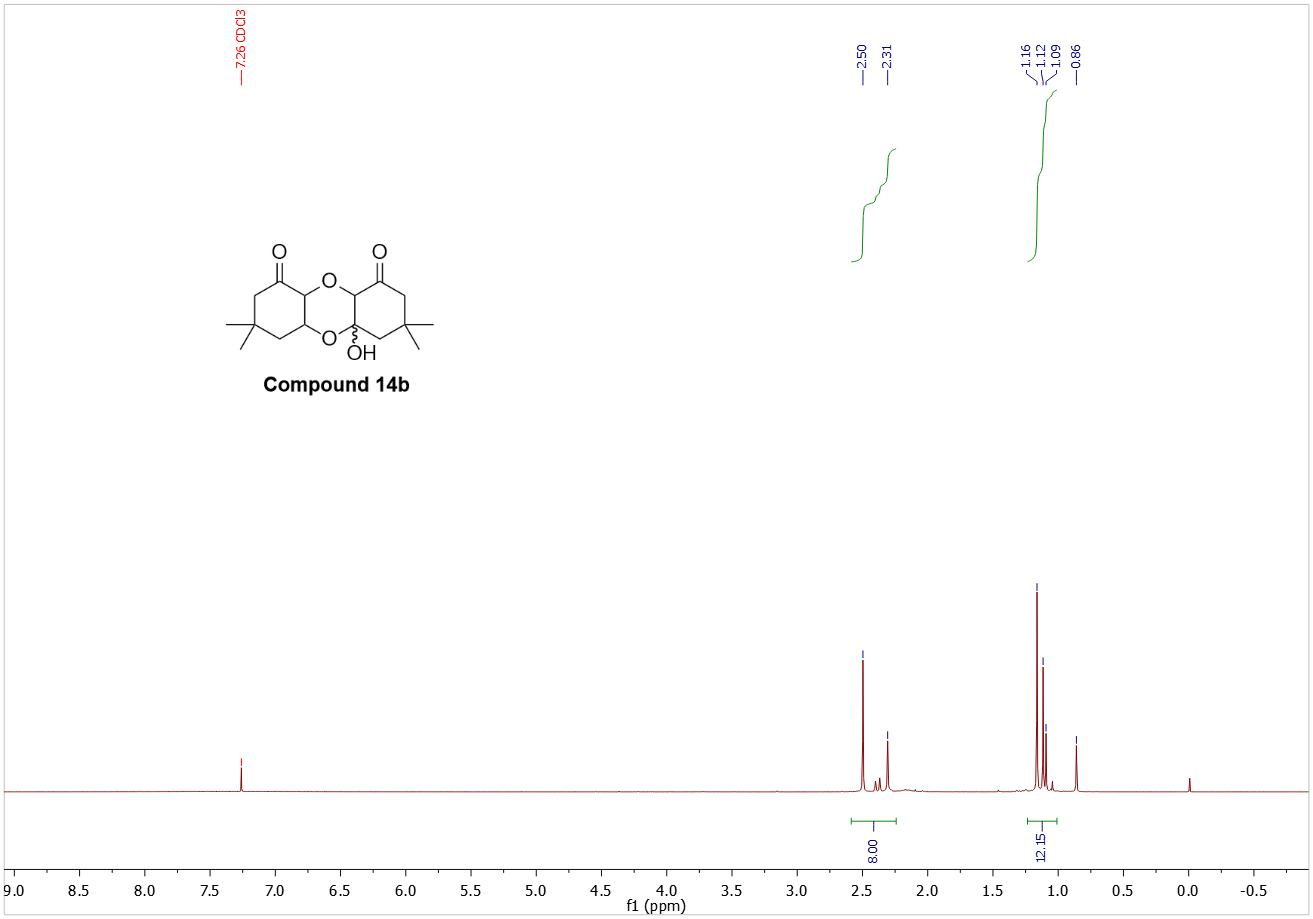

**^1^H NMR of compound 14 (in an equilibrium favoring 14b in CDCl_3_)**

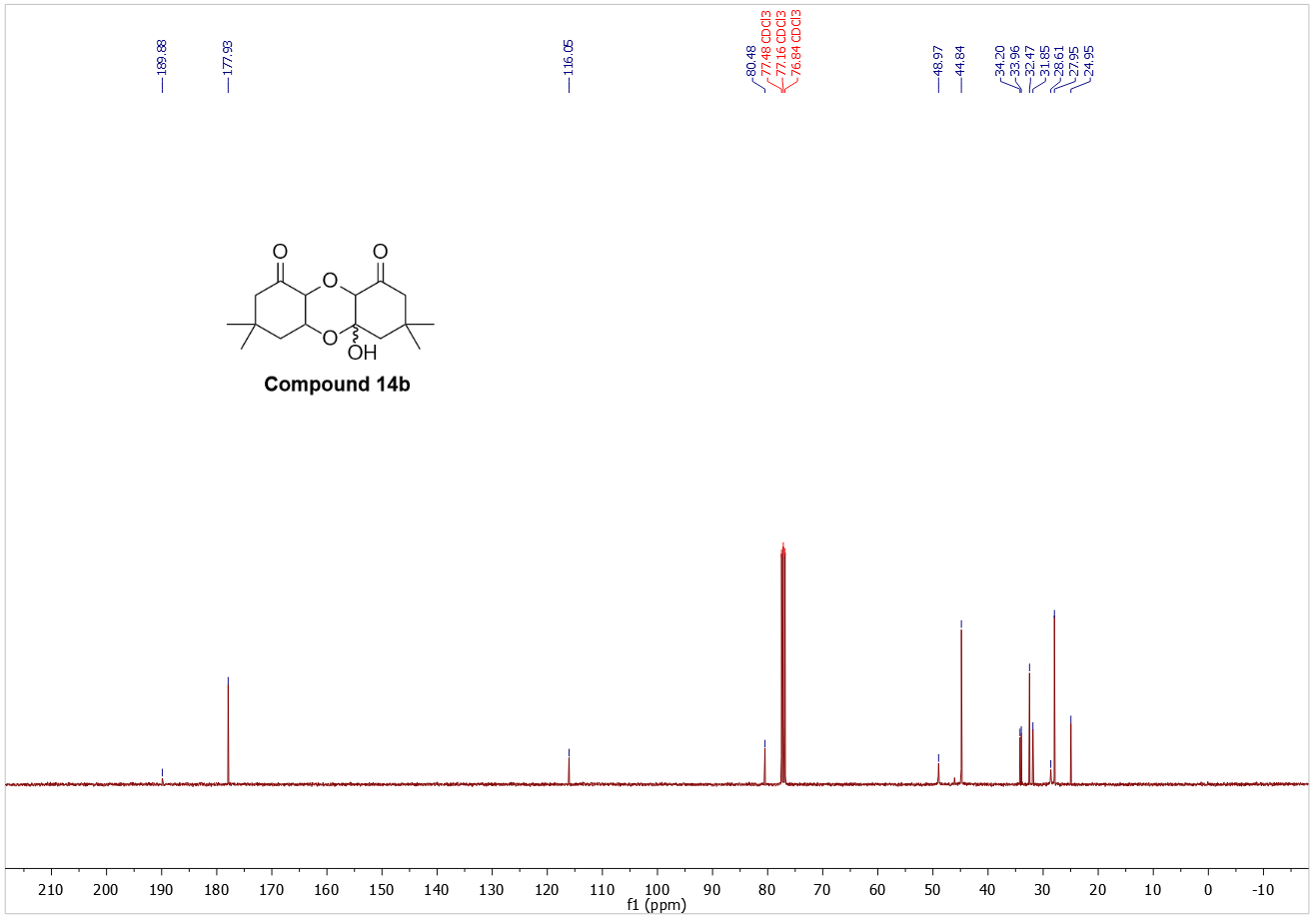

**^13^C NMR of compound 14 (in an equilibrium favoring 14b in CDCl_3_)**
